# Lactate dehydrogenase is the Achilles’ heel of Lyme disease bacterium *Borreliella burgdorferi*

**DOI:** 10.1101/2025.02.07.637162

**Authors:** Ching Wooen Sze, Michael J Lynch, Kai Zhang, David B. Neau, Steven E. Ealick, Brian R Crane, Chunhao Li

## Abstract

As a zoonotic pathogen, the Lyme disease bacterium *Borreliella burgdorferi* has evolved unique metabolic pathways, some of which are specific and essential for its survival and thus present as ideal targets for developing new therapeutics. *B. burgdorferi* dispenses with the use of thiamin as a cofactor and relies on lactate dehydrogenase (BbLDH) to convert pyruvate to lactate for balancing NADH/NAD^+^ ratios. This report first demonstrates that BbLDH is a canonical LDH with some unique biochemical and structural features. A loss-of-function study then reveals that BbLDH is essential for *B. burgdorferi* survival and infectivity, highlighting its therapeutic potential. Drug screening identifies four previously unknown LDH inhibitors with minimal cytotoxicity, two of which inhibit *B. burgdorferi* growth. This study provides mechanistic insights into the function of BbLDH in the pathophysiology of *B. burgdorferi* and lays the groundwork for developing genus-specific metabolic inhibitors against *B. burgdorferi* and potentially other tick-borne pathogens as well.

## Introduction

Lyme disease (LD) is an emerging vector-borne illness caused by *Borreliella* (formerly known as *Borrelia*) *burgdorferi*(1). Currently, LD is most commonly reported in the Europe and North America but some cases have also been reported in Asia and South America(2). Between year 2010-2018, approximately 476,000 new LD cases were reported annually in the U.S. alone(3–5). There is no current vaccine for LD; vaccines based on the outer surface protein A (OspA) were withdrawn or discontinued owing to safety concerns(6–8). Although an OspC-based vaccine is available for dogs, an analogous vaccine has not yet been developed for humans(6, 9, 10). Antibiotics treat LD effectively if administered soon after infection; however, long-term LD has proven resistant to treatment and the ability of common therapies to eliminate the infection has been called into question(11, 12). In Post-Treatment Lyme Disease Syndrome (PTLDS) at least 25% of patients continue to have joint inflammation after treatment with antibiotics and over 10% of patients who receive early treatment develop long-term symptoms such as fatigue, cognitive deficits, and musculoskeletal pain(11–13). Moreover, PTLDS patients often fail to respond to further antibiotic treatments(13, 14). Such enduring symptoms may result from *B. burgdorferi* forming drug-resistant “persistent” cells(15, 16). Indeed, in the presence of the routinely administered antibiotics such as doxycycline, amoxicillin, and ceftriaxone, *B. burgdorferi* adapts to a slow growing state(17–20), which is similar to that found within the nutrient-poor environment of the tick. Therefore, there is an emerging demand for development of novel therapeutics to treat LD or as a prophylactic to prevent LD(14).

The genome of *B. burgdorferi* consists of a ∼910 kilobases linear chromosome and more than 20 circular and linear plasmids(21, 22). With such a small genome, *B. burgdorferi* has a minimal metabolic system and thus relies heavily on tick vectors and mammalian hosts to provide constitutive nutrients. One such example is how *B. burgdorferi* lacks genes that encode the proteins necessary for lipid, nucleotide, amino acid, and cofactor synthesis. Instead, its genome contains 16 genes for broad-spectrum membrane transporters presumably responsible for up-taking these essential nutrients from the extracellular environment(21). Intriguingly, *B. burgdorferi* also dispenses with the use of thiamin(23), which was thought to be essential for all living organisms due to its vital role as cofactor in central carbohydrate metabolism and amino acid biosynthesis(24, 25). No biosynthesis pathways, transporters, or enzymes that use thiamin as a cofactor can be found in the genome of *B. burgdorferi*, which is unprecedented(23). In addition, *B. burgdorferi* also lacks genes that encode enzymes involved in the tricarboxylic acid (TCA) cycle(21), e.g., pyruvate dehydrogenase (PDH) that decarboxylates pyruvate producing acetyl-CoA and CO_2_. Pyruvate is a central metabolite for the generation of cofactors and energy needed for the anabolic and catabolic reactions of the cell(26). Moreover, pyruvate functions as a protective molecule for *B. burgdorferi* against reactive oxygen species (ROS) that cause DNA damage and killing(27). To compensate for the lack of thiamin, PDH, and the TCA cycle, *Borreliella* species have evolved a distinct metabolic alternative that relies on lactate dehydrogenase (LDH) for continuous ATP synthesis and the regeneration of NADH/NAD^+^ through anaerobic glycolysis(21, 28–30).

The LDH of *B. burgdorferi* (BbLDH) is encoded by the gene *bb_0087*(21). LDH is the final enzyme in anaerobic glycolysis which is responsible for replenishing healthy NADH/NAD^+^ ratios through the conversion of pyruvate to lactate and vice versa(31, 32). Maintaining a balance between NADH and NAD^+^ is of the utmost importance for cells as their relative amounts reflect the intracellular redox state. Unlike other bacteria that can utilize tryptophan or aspartic acid for NAD^+^ regeneration(33, 34), *B. burgdorferi* relies on glycolysis to regenerate NADH and lactogenesis to replenish NAD^+^(21). In addition to upholding a balanced intracellular pool of NAD, LDH is also essential for maintaining the pyruvate level in *B. burgdorferi* by reverting lactate into pyruvate(30). There are two major subcategories of LDH enzymes, L(+)-LDHs and D(-)-LDHs, which are distinguished by the stereoisomeric form of lactate they produce(30). BbLDH is L(+) whereas human LDH (HsLDH) is L(-). The L(+)-LDHs are typically homotetrameric and are frequently allosterically regulated, most commonly by fructose-1,6-bisphosphate (FBP)(35, 36). These enzymes typically exhibit sigmoidal kinetic profiles unless in the presence of an allosteric binding partner, which is required for high catalytic activity(30). Studies have shown that the sensitivity of LDH to FBP varies across species of bacteria due to the FBP binding site(37). Furthermore, phosphate ions can act as inhibitors or activators of LDHs in different bacteria species depending on the presence or absence of FBP and the pH of the surrounding environment(37). All of these factors suggest a complex interplay between cofactors and environmental factors in the regulation of bacterial LDH.

BbLDH has not yet been functionally or structurally characterized and its importance in *B. burgdorferi* survival and infectivity has not been established. In this report, we decipher the biochemical and structural feature of BbLDH, establish its role in the pathophysiology of *B. burgdorferi*, and then explore its potential as a new therapeutic target to treat Lyme disease.

## Results

### BB_0087 has LDH activity

BB_0087 (BbLDH) comprises 316 residues and is annotated as the sole lactate dehydrogenase (LDH) in the genome of *B. burgdorferi*(21). It is well conserved among all sequenced *Borrelia* species with a ∼82% shared sequence identity (Extended Data Fig. 1). To establish the LDH activity of BB_0087, we expressed and purified a full-length N-terminal His- and FLAG-tagged BB_0087 recombinant protein (rBB_0087) under native conditions using affinity and size exclusion chromatography (SEC) (Extended Data Fig. 2A). The ability of rBB_0087 to reduce NAD to NADH was quantified spectrophotometrically using the Lactate Dehydrogenase Activity Assay Kit. The result showed that BB_0087 can generate NADH from lactate (Extended Data Fig. 2B), which was completely abolished by gossypol, a non-selective LDH inhibitor(38–41). In addition, site-directed mutations in three conserved residues found in LDH enzymes (Thr146, Arg154, and His178) significantly reduced if not abolished the NADH-dependent activity of BB_0087 (Extended Data Fig. 2B, C). These results indicate that BB_0087 is a genuine LDH with conserved catalytic residues commonly found in other LDHs that can be targeted by LDH inhibitor.

### BbLDH is allosterically regulated by fructose 1,6-bisphosphate

Unlike eukaryotic LDH, some bacterial LDHs require an allosteric regulator, fructose 1,6-bisphosphate (FBP), to achieve optimum activity(30, 42–44). The allosteric regulation of bacterial LDH was first elucidated from the structural analyses of *Bacillus stearothermophilus* LDH (BsLDH)(45). FBP was discovered to interact with Arg173 and His188, two positively charged residues in each of the two juxtaposed subunit, thereby enhancing the BsLDH tetramer formation and catalytic activity. Sequence alignment showed that BbLDH also contains these two conserved residues (Arg156 and His171) at the same position found in BsLDH (Extended Data Fig. 3), suggesting a common regulatory mechanism. To confirm this, the LDH assay was performed with and without the addition of 3 mM FBP. We found that addition of FBP decreased the pyruvate *K*_half_ of BB_0087 from 6.2 mM (with a 95% confidence range [95%CI] of 3 – 16 mM) to 4.3 mM (95% CI: 2 – 12 mM) (**Fig. 1A**). In addition, mutation of the key residue His171 to Ala completely abolished the FBP-mediated allosteric activation on BB_0087 (**Fig. 1A**) as His171 forms eight hydrogen bonds with FBP (four on each chain) to stabilize the interaction between BbLDH and FBP (**Fig. 2D**). This result confirms that BB_0087 is allosterically regulated via FBP.

**Figure 1.**
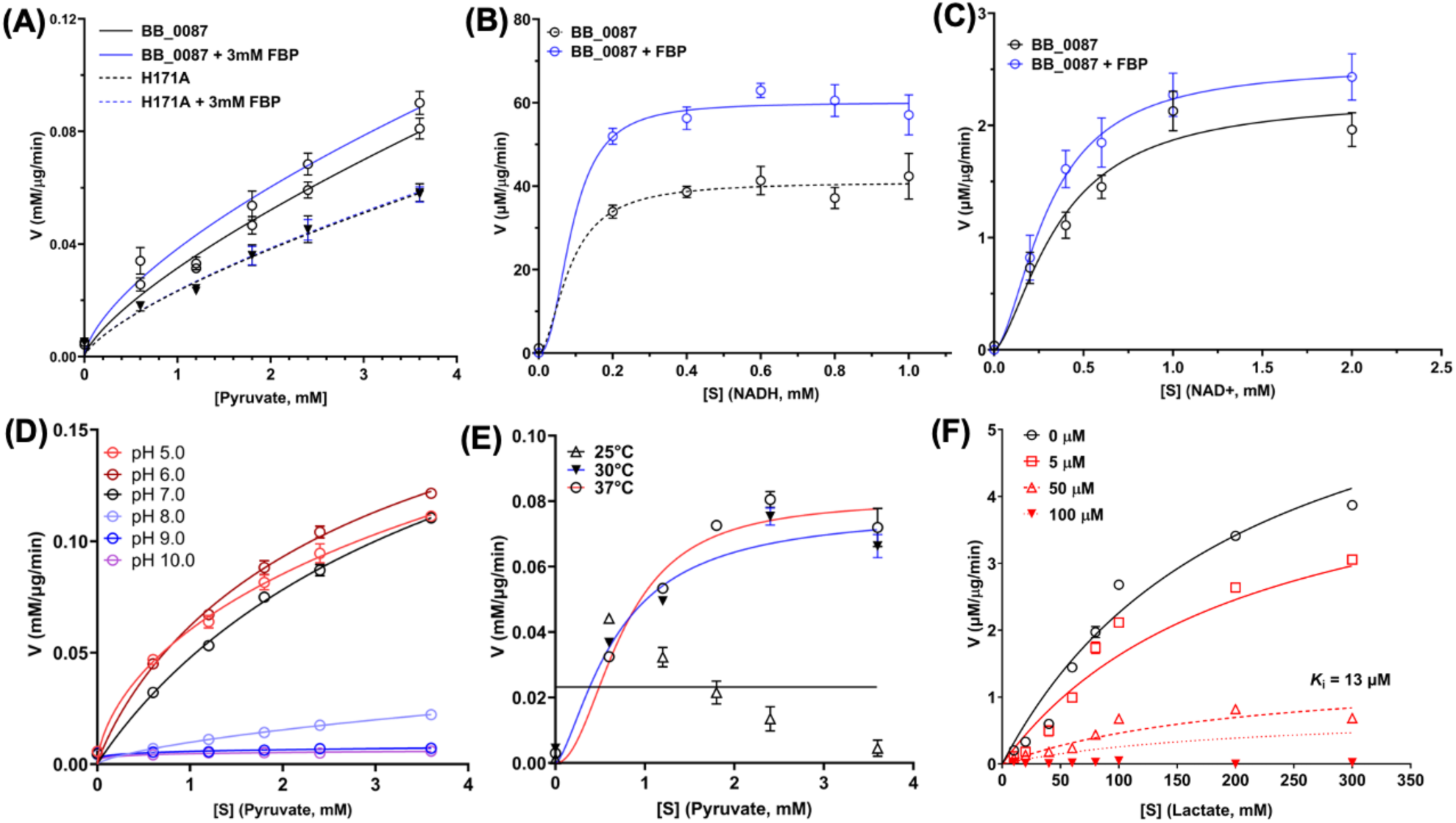
BbLDH enzyme kinetics and inhibition constants of gossypol against BbLDH. **(A)** BB_0087 is allosterically regulated by fructose-1,6-bisphosphate (FBP). Addition of 3 mM FBP enhanced the binding affinity of BbLDH to pyruvate. Mutation of H171A, a predicted key residue in the FBP binding site, abolished the allosteric activation of FBP on BbLDH. **(B & C)** Addition of FBP increased the *V*_max_ of BbLDH for NADH and NAD^+^ using a varying concentration of cofactors. **(D)** BbLDH has an optimal working pH range between 5.0 to 7.0. **(E)** BbLDH has an optimal working temperature of 37°C using a varying concentrations of pyruvate substrate. **(F)** Enzyme kinetics of inhibition on BbLDH by gossypol. BB_0087 catalyzed the reaction of lactate to pyruvate which could be inhibited by gossypol in a dose-dependent manner. Of note, NADH standards of 0 (blank), 2.5, 5, 7.5, 10, and 12.5 nmol per well were generated using the spectrophotometric assay at OD_340_. The linear regression and enzyme kinetic analyses were performed in GraphPad Prism 10. S, substrate, V, velocity.

**Figure 2:**
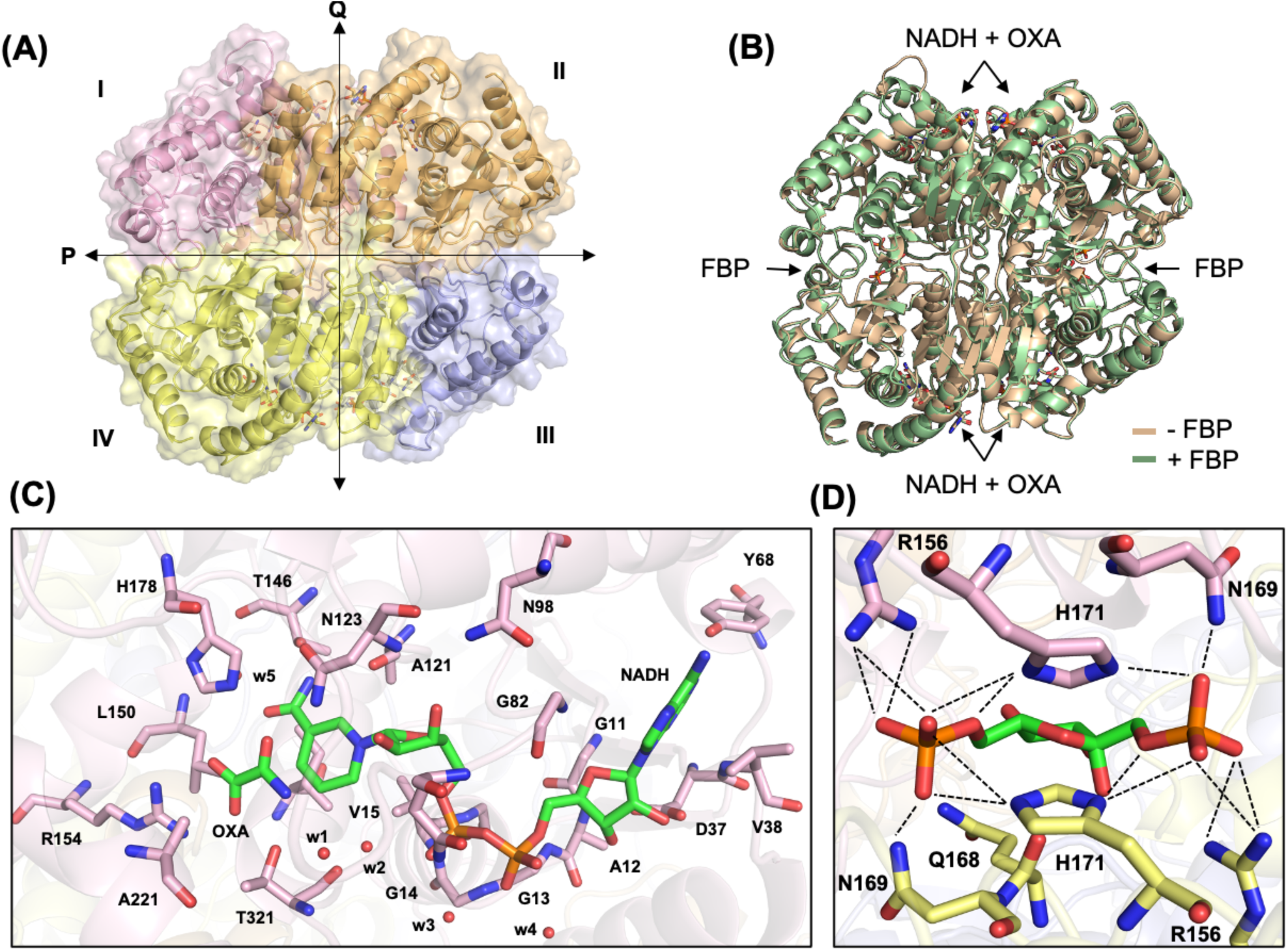
Structure of *B. burgdorferi* LDH (BbLDH). **(A)** 2.1 Å resolution crystal structure of BbLDH tetramer with NADH and oxamate (PDB 9DQ9). **(B)** Superimposition of BbLDH crystal structure without FBP (tan, PDB 9DQ9) and with FBP (green, PDB 9DQ7) (RMSD = 0.230 Å, 314 atom pairs). **(C)** Interacting residues in the NADH and oxamate binding site pocket. **(D)** Salt bridge and hydrogen bonding residues that interact with FBP in BbLDH (chain A and C, pink and yellow).

### Characterization of BbLDH activity

To further characterize the activity of BbLDH, the enzyme kinetics for NADH, lactate, and NAD^+^ were measured with or without the addition of 3 mM FBP. The results showed that BbLDH can catalyze the reversible conversion between pyruvate and lactate (**Fig. 1A**). Addition of 3 mM FBP significantly increases the *V*_max_ for NADH from 41 to 60 µM/µg/min, and *V*_max_ for NAD^+^ from 2.2 (95% CI: 1.9 - 3.2) to 2.5 (95% CI: 2.1 – 4.1) µM/µg/min (**Fig. 1B, C**). The optimal pH for the activity of BbLDH was between pH 5.0 – 7.0 (**Fig. 1D**), and the optimal working temperature was at 37°C (**Fig. 1E**). We further evaluated the inhibitory effect of gossypol on the activity of BbLDH with respect to NADH reduction and oxidation and the results showed a dose-dependent inhibition of gossypol on the LDH activity of BbLDH with a *K*_i_ of 13 µM (95% CI: 11 – 14 µM) (**Fig. 1F**).

### Structure of BbLDH with and without FBP

The crystal structures of BbLDH with and without FBP were determined via X-ray crystallography to a resolution of 2.1 Å (**Fig. 2A** & **B**). Both structures were determined in the presence of NADH and the pyruvate analog oxamate (OXA). BbLDH forms a homo-tetramer or a dimer of dimers, wherein each monomer comprises eight a-helices and five β-strands (**Fig. 2A**) and contains binding sites for oxamate, NADH, and FBP (**Fig. 2B** and Extended Data Fig. 4). Interestingly, FBP binding does not change the overall structure of the BbLDH tetramer to any appreciable extent, with the FBP-bound tetramer superimposing on the apo-tetramer with an RMSD of 0.230 Å (**Fig. 2B**). BbLDH also shares a high degree of structural similarity to the dimeric human LDH (HsLDH, PDB: 8FW6, Extended Data Fig. 4A) as well as the tetrameric BsLDH (PDB: 1LDB, Extended Data Fig. 4B). Within the BbLDH tetramer, the four NADH/oxamate binding sites (one per subunit) are highly similar, as are the two FBP binding sites (one per dimer) (Extended Data Fig. 4C,D), with the only major difference being the relative position of the Gln85-Arg91 loop adjacent to the NADH/oxamate binding site (Extended Data Fig. 4D). Within each binding site, several conserved residues interact directly with NAD/oxamate (**Fig. 2C**) and FBP (**Fig. 2D**). LDH tetramers assume either an R- or T-state distinguished largely by the relative positioning of the two dimers and depending on the binding of substrate and allosteric regulators. These subunit states influence a loop and helix conformation that borders the catalytic site(36, 42). The BbLDH subunits show some variability in the loop configuration but are generally similar to one another. The subunits appear primarily in the R-state, which is consistent with the binding of oxamate (Extended Data Fig. 5). Details of the interactions between the enzyme and oxamate, NAD, or FBP are also tabulated in Extended Data Tables 2 to 4. Although BbLDH crystalizes as a tetramer, SEC-MALS of BbLDH in solution indicates that BbLDH is predominantly a dimer in solution at concentrations of 10 μM – 20 μM, regardless of the presence of NADH, oxamate, FBP, or any combination of the three compounds (Extended Data Fig. 6).

### BbLDH is required for *B. burgdorferi in vitro* growth

To study the function and significance of BbLDH in the life cycle of *B. burgdorferi*, we attempted to construct a complete knockout of *bb_0087* but were unsuccessful after multiple attempts, likely because the deletion of this gene was lethal to the spirochetes. To overcome this issue, we generated an IPTG-inducible conditional knockout strain of *bb_0087* (87^mut^) in the infectious B31 A3-68 (WT) strain(46), as previously reported(47). In brief, the intact *bb_0087* gene was cloned into pJSB275(47), an IPTG-inducible shuttle vector (**Fig. 3A**). The resultant vector was transformed into the WT which was then used as the parental strain to in-frame replace the *bb_0087* gene on the chromosome with *kan*, a kanamycin-resistant marker (**Fig. 3B**). To monitor the expression of *bb_0087*, a FLAG tag was added to the C-terminal of BB_0087. *In vitro* growth analysis at 34°C detected no BbLDH protein in 87^mut^ (**Fig. 3C**) and the mutant was unable to grow without IPTG (**Fig. 3D**). The addition of 1 mM IPTG successfully restored the expression of BbLDH in 87^mut^ (**Fig. 3C**) and its growth rate to the wild-type level (**Fig. 3D**). Collectively, these results indicate that BbLDH is vital for *B. burgdorferi* growth *in vitro*, making it a viable candidate for drug target candidate.

**Figure 3.**
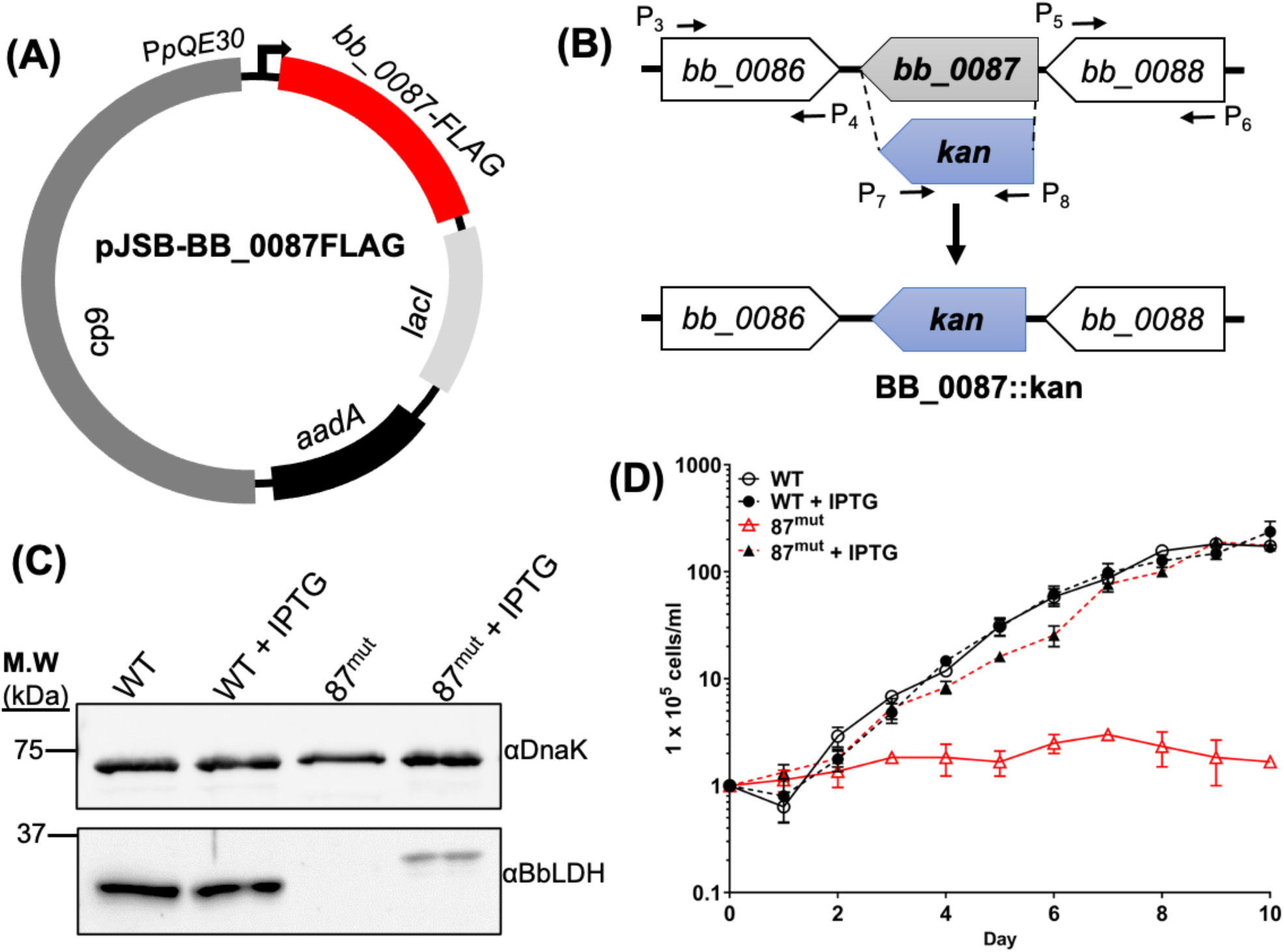
BB_0087 is essential for the *in vitro* growth of *B. burgdorferi*. **(A)** To create an inducible vector for condition knockout of *bb_0087*, the full-length *bb_0087* gene was cloned into an IPTG-inducible vector (47) with a C-terminal FLAG tag, generating pJSB-BB_0087FLAG. The final construct was transformed into the wild-type B31 A3-68 parental strain. **(B)** To construct an *in-frame* deletion mutant in WT carrying pJSB-BB_0087FLAG, the primer pairs P_3_/P_4_ and P_5_/P_6_ were used to amplify the upstream and downstream flanking regions of *bb_0087*. The primer pair P_7_/P_8_ was used to amplify a promoterless kanamycin resistance cassette (*kan*). *bb_0087 orf* was in-frame replaced by *kan* via the PCR fusion technique with the primer pair P_3_/P_6_. The resulting PCR amplicon was cloned into a pGEM-T-easy vector forming BB_0087::kan. **(C)** Immunoblot analysis of WT and 87^mut^ in the presence or absence of IPTG from the growth curve in **(D)** at day 10. The same amounts of WT and 87^mut^ whole-cell lysates were analyzed by SDS-PAGE and then probed with antibodies against BbLDH and DnaK. Of note, the band detected in the 87^mut^ strain was larger than its native form in the WT due to the addition of FLAG tag. **(D)** Growth analysis of WT and 87^mut^ under routine laboratory condition (34°C/pH 7.4) with and without 1 mM IPTG addition to induce the expression of BbLDH. Cell numbers were enumerated daily until cells entered the stationary growth phase (∼10^8^ cells/ml). Cell counting was repeated in triplicate with three independent samples, and the results are expressed as means ± SEM.

### BbLDH contributes to the infectivity of *B. burgdorferi* in mice

Our growth analysis indicated that BbLDH is absolutely essential for the growth of *B. burgdorferi in vitro*. To examine the fitness of this mutant as well as the significance of BbLDH in the pathogenesis of *B. burgdorferi*, a mouse infection study was conducted using WT and 87^mut^ with or without supplementation of IPTG in the drinking water to induce the expression of BbLDH in mice. Three weeks after the infection, mice were sacrificed. Ear tissues were harvested for qRT-PCR and blood samples were used for seroconversion analysis. qRT-PCR data showed the presence of *flaB* mRNA in the ear tissues in both the WT and 87^mut^ infected mice, indicating that the mutant was able to disseminate from the initial inoculation site to distal organs. However, mice receiving IPTG in their drinking water showed a five-fold higher level of *flaB* transcript as compared to those that did not (**Fig. 4A**), suggesting that IPTG induction of BB_0087 could enhance the fitness and replication of the 87^mut^ *in vivo*. In addition, the mice receiving IPTG-supplemented water showed sign of seroconversion (4/4) while the group that did not receive IPTG had only one mouse with a weak sign of seroconversion (1/4) (**Fig. 4B**). These results indicate that BbLDH is essential for the infectivity and fitness of *B. burgdorferi* in mammalian hosts.

**Figure 4.**
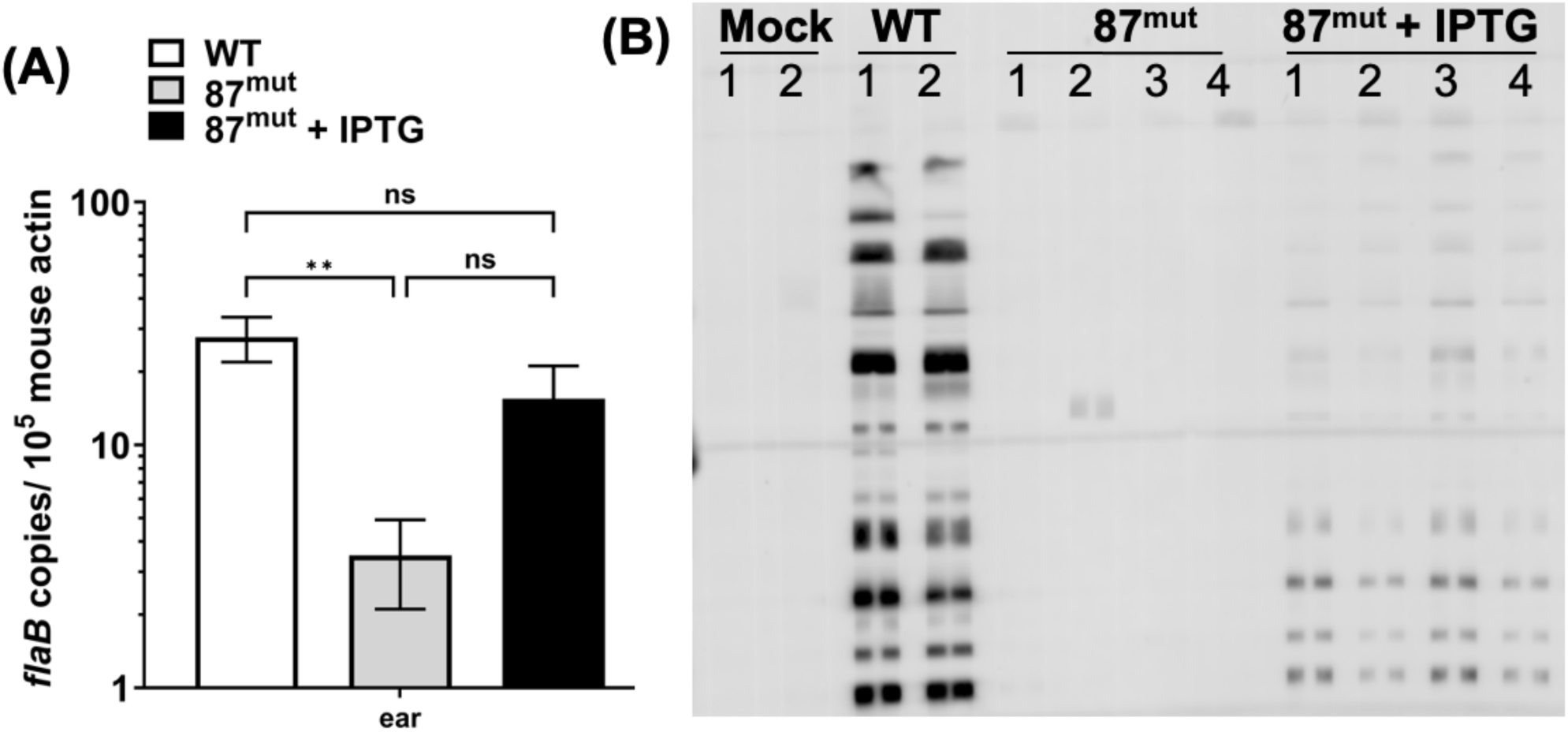
BbLDH contributes to the infectivity of *B. burgdorferi.* To determine the role of BbLDH in the pathophysiology of *B. burgdorferi*, a needle infection study was conducted using BALB/c mice. 10^3^ of WT or 87^mut^ strains were subcutaneously inoculated into BALB/c mice and sacrificed three weeks after injection. Mice infected with 87^mut^ were divided into two groups (4 mice per group), one group received regular drinking water while the other received IPTG supplemented water as described(93) for the duration of the infection study. **(A)** Ear tissues were harvested for qRT-PCR as described(92) and **(B)** the blood was collected for seroconversion test. Each lane represents serum obtained from individual mice under each group as indicated on the top of the image. qRT-PCR data are presented as mean *flaB* transcript over 10^5^ of mouse *β-actin* ± SEM, significant difference (\**P* < 0.05; ** *P* < 0.01). ns: no significance.

### Identification of novel LDH inhibitors via small compound library screening

Given the significance of BbLDH in the metabolism and pathogenicity of *B. burgdorferi*, it represents as a viable target for developing specific metabolic inhibitors against LD. A recent study by Lynch *et al.*(48) as well as data from this study indicated that *B. burgdorferi* is susceptible to inhibition by commercially available broad spectrum LDH inhibitors. To discover additional LDH inhibitors, we first screened the Natural Products Set IV library (NPS-IV, containing 419 compounds) from the National Cancer Institute (NCI). Using a fluorescence-based high-throughput screening (HTS) assay(49), 23 compounds showed inhibitory effect on the LDH activity of BbLDH when compared to DMSO-treated wells at 10 μM concentration during the first screening (Extended Data Fig. 6A). A follow-up secondary screening further confirmed the anti-LDH activity of four lead compounds (**Fig. 5A**) with inhibitory effect on the LDH activity of BbLDH in the µM range (**Fig. 5C-F**). These four lead compounds are structurally diverse from gossypol (**Fig. 5A**). In comparison to gossypol which has a *K*_i_ of 36 µM (**Fig. 5B**, 95%CI:16 to 86 µM), compound 45923 showed the most potent and promising inhibitory effect on BbLDH, with a *K*_i_ value of 54 µM (**Fig. 5D**, 95%CI: 30 to 103 µM), followed by 14975 (**Fig. 5C**, *K*_i_ = 66 µM, 95%CI: 38 to 123 µM), 350085 (**Fig. 5F**, *K*_i_ = 205 µM, 95%CI:143 to 319 µM), and 114344 (**Fig. 5E**, *K*_i_ = 349 µM, 95%CI: 217 to 701 µM). Among these four compounds, 45923 is the smallest molecule with a molecular weight of 216.0 Dalton (**Fig. 5A**).

**Figure 5.**
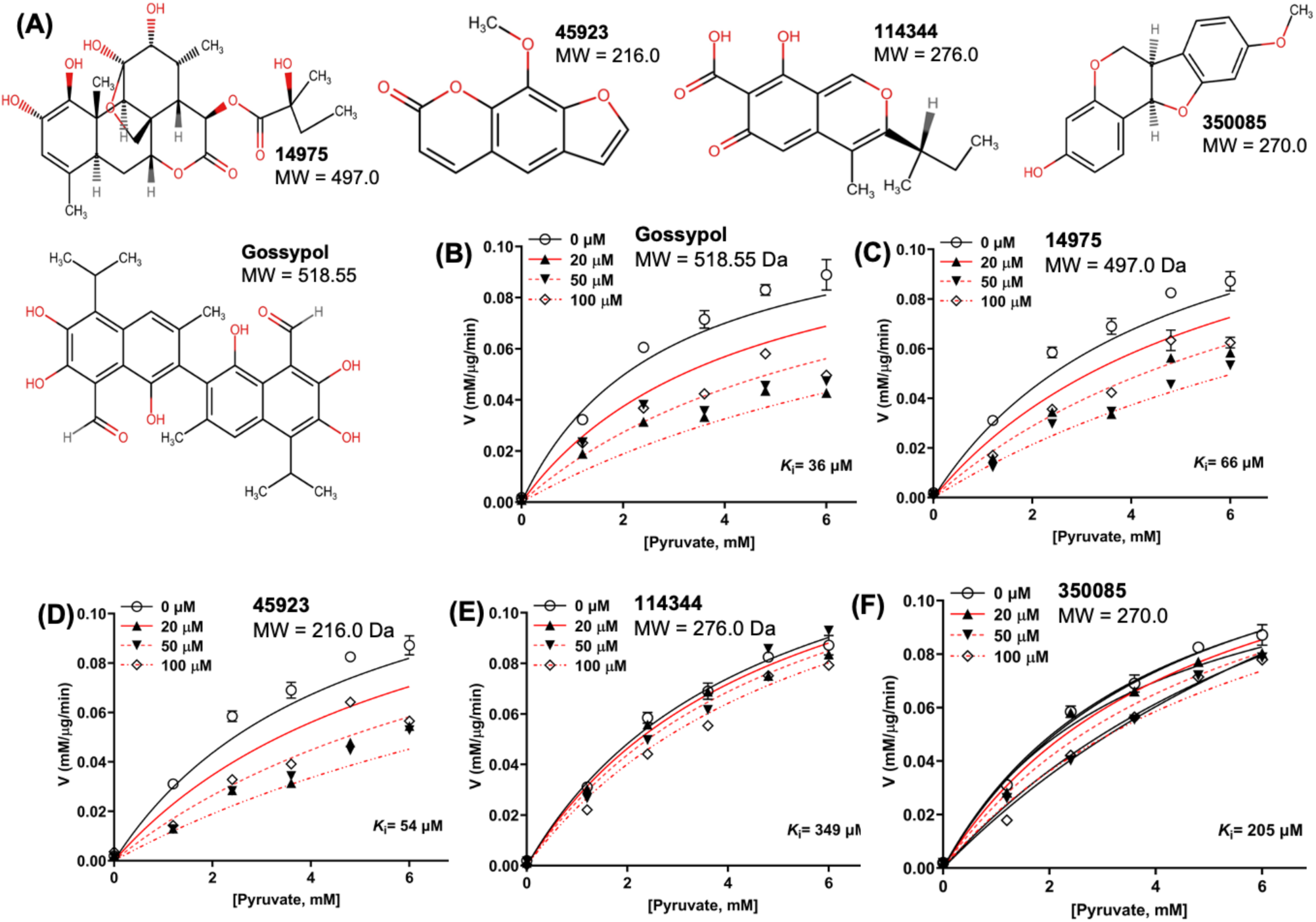
BbLDH activity can be inhibited by gossypol and four novel LDH inhibitors. Four lead compounds with anti-LDH activities were identified through high-throughput screening from the Natural Product Set IV library. **(A)** Chemical structures of the four lead compounds with inhibitory effect on BbLDH activity along with their molecular weight. Gossypol was included for comparison. Enzyme kinetics of inhibition of BbLDH by 0 – 100 µM of **(B)** gossypol; **(C)** 14975; **(D)** 45923; **(E)** 114344; and **(F)** 350085 using varying concentrations of pyruvate as a substrate. Data are presented as mean velocity ± SEM from two independent experiments.

### Small molecule inhibitors can inhibit the growth of *B. burgdorferi in vitro*

The four newly identified LDH inhibitors showed varying degrees of inhibition on BbLDH activity (**Fig. 5C-F**). To determine the inhibitory effect of LDH inhibitors on the growth of *B. burgdorferi*, we performed *in vitro* growth analysis using various concentration of inhibitors between 0 – 300 µM with gossypol and DMSO serving as positive and negative controls, respectively. Gossypol showed inhibitory impact on the growth of *B. burgdorferi* at 100 µM (**Fig. 6A**). Among the four newly identified compounds, 45923 showed the highest potency with inhibitory effect observed at 100 µM, which was comparable to gossypol (**Fig. 6C**). This result is consistent with 45923 showing the lowest *K*_i_ for BbLDH among the four small molecules (**Fig. 5D**). At 200 µM and 300 µM concentration, 45923 inhibited the growth of *B. burgdorferi* between 99 – 100%, similar to what was observed using gossypol (**Fig. 6A, C**). The remaining three molecules only exhibited ∼ 50% growth inhibition at 200 – 300 µM concentrations (**Fig. 6B, D, E**). Combining all inhibitor analyses, 45923 is the most promising candidate as a novel LDH inhibitor.

**Figure 6.**
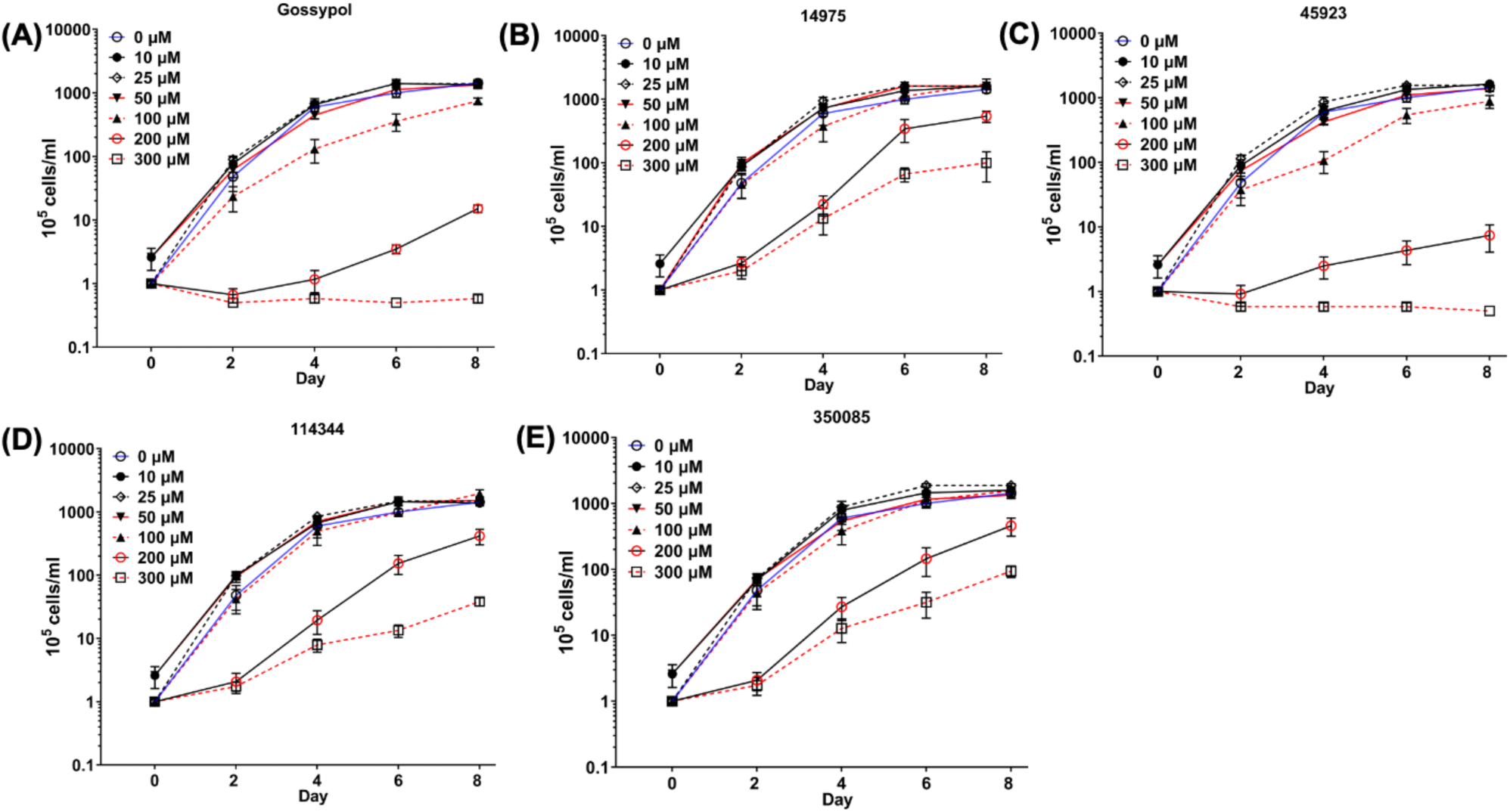
Impact of LDH inhibitors on the growth of *B. burgdorferi.* Growth analysis of wild-type *B. burgdorferi* in the presence of 0 (DMSO) to 300 µM of **(A)** gossypol; **(B)** 14975; **(C)** 45923; **(D)** 114344; or **(E)** 350085. Cell numbers were enumerated every two days using a Petroff-Hausser counting chamber until cells entered the stationary growth phase (∼10^8^ cells/ml). Cell counting was repeated in triplicate with two independent samples, and the results are expressed as means ± SEM.

### Inhibitor docking analysis of BbLDH

Our attempts to crystallize BbLDH with gossypol or any of the four inhibitors identified in our HTS assays failed to yield any structures of inhibitor-bound LDH regardless of the presence or absence of NADH, oxamate or FBP. Therefore, to identify how each inhibitor interacts with BbLDH, we docked each compound to LDH *in silico* using the Schrodinger software suite. We first generated either an apo-LDH model that lacked NADH and oxamate, or a cofactor-bound LDH model (**Fig. 7A**). Lineweaver–Burk plots for gossypol, 45923, 14975, 114344 and 350085 showed that gossypol exhibits uncompetitive inhibition for pyruvate substrate, while 45923, 14975, 114344 and 350085 were competitive inhibitors (Extended Data Fig. 7). Therefore, gossypol was docked to our BbLDH model that contained NADH and oxamate, while the other four inhibitors were docked to the apo LDH model. To lend support to our docking results of compounds 45923, 14975, 114344 and 350085, we compared the top docking pose of gossypol (docking score = -3.889) to the recent crystal structure of gossypol in complex with HsLDH(50) (**Fig. 7** & Extended Data Fig. 8). Overall, gossypol is predicted to bind to BbLDH in a manner similar to how it binds to HsLDH by interacting with residues in a loop adjacent to the NADH and lactate/oxamate binding pockets (Extended Data Fig. 8). In HsLDH, gossypol participates in hydrogen bonding and salt bridge interactions with Asn110 and Arg111. In BbLDH, Asn110 and Arg111 correspond to Asp96 and Lys97, which also interact with gossypol. Similarly, in HsLDH, residue Glu103 forms a hydrogen bond with the aldehyde carbonyl oxygen of gossypol. In BbLDH, the equivalent residue, Glu89, is in proximity (4.3 Å) to the analogous aldehyde carbonyl oxygen. Analysis of our docking results also revealed a hydrogen bond between Lys86 and a hydroxyl group ortho to the aldehyde group in gossypol.

**Figure 7.**
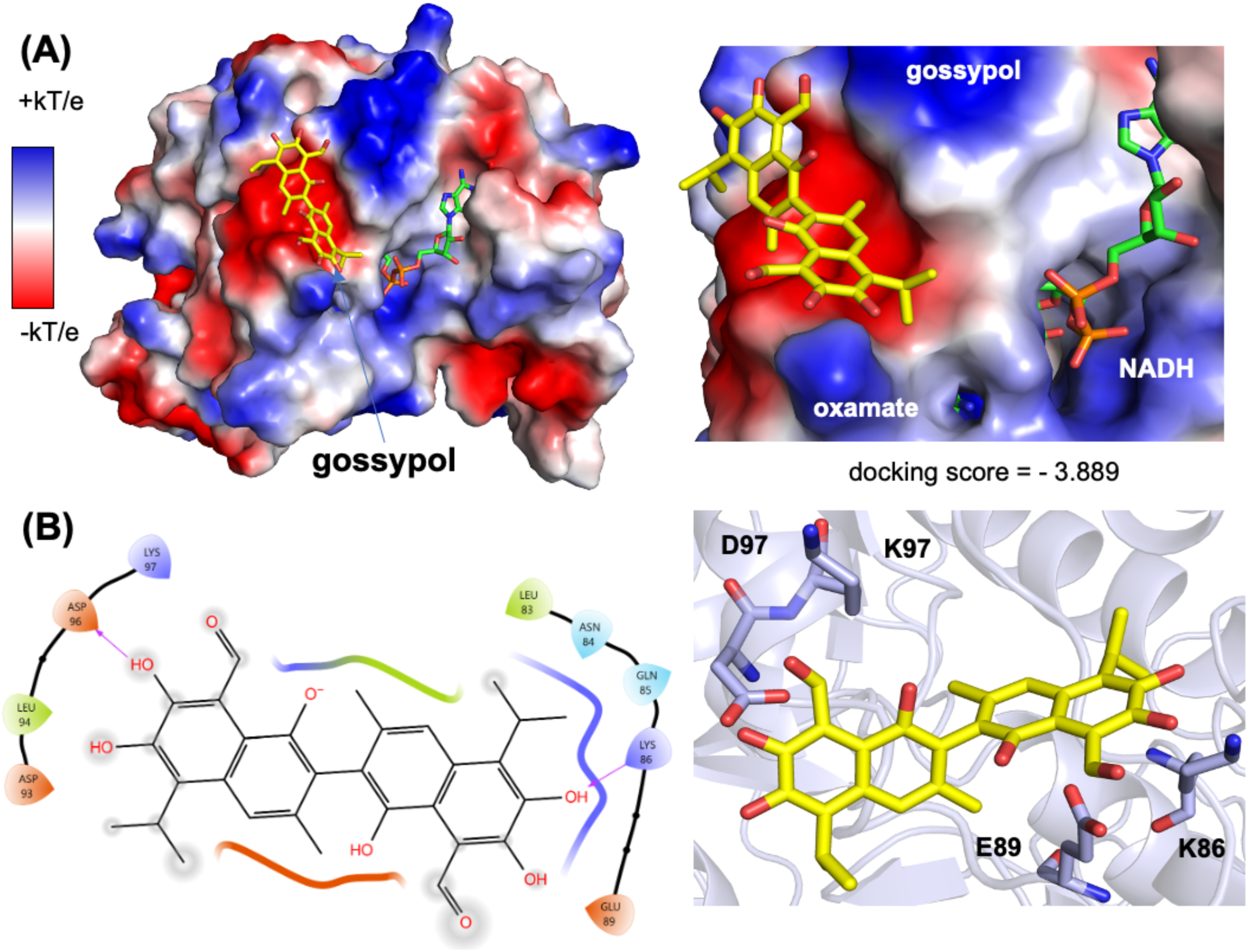
Docking analysis of gossypol. **(A)** Top docking pose of gossypol to an NADH- and oxamate-bound BbLDH model. (left) Overview of BbLDH dimer model with electrostatic potential surface. (right) The gossypol binding site location relative to the NADH and oxamate cofactors. **(B)** Hydrogen bonding interactions between gossypol and BbLDH. A ligand interaction diagram (left) and stick model (right) for the top-ranked gossypol pose are shown.

Using a similar approach, we docked 45923, 14975, 114344 and 350085 to the apo-BbLDH model (Extended Data Figs. 9, 10). Overall, all four inhibitors localize to the NADH and lactate binding pockets, consistent with their competitive inhibition with pyruvate (Extended Data Fig. 7). Each inhibitor is predicted to interact with a series of residues within the binding pocket, including: Gly13, Gly14, Val15, Gly16, Asn84, Asp37 and Thr80 (Extended Data Fig. 10). In total, 14975 and 114344 form four hydrogen bonding contacts with BbLDH, 45923 forms a single hydrogen bond with LDH and 350085 forms no hydrogen bonds or salt-bridge contacts with Bb LDH. For 45923 and 350085, the change in solvent-accessible hydrophobic surface area of the compounds upon binding may be an important determinant in binding affinity.

### Small molecule inhibitors inhibit the growth of cancer cell *in vitro*

Of the four newly identified LDH inhibitors, 45923 (Methoxsalen) and 350085 (Medicarpin) are commercially available. To determine the inhibitory effect of these two compounds on eukaryotic cells, a growth inhibition study was performed using a well-established cancer cell line, HeLa cells. DMSO was included as negative control while gossypol was used as positive control. A non-cancerous cell line, Telomerase Immortalized Gingival Keratinocytes (TIGKs),(51) was included to determine the toxicity of the inhibitors to normal human cells. Gossypol was highly inhibitory to both HeLa and TIGKs cells at as low as 1 – 2 µM concentration (**Fig. 8A**). 45923 began exhibiting significant growth inhibition on HeLa cells at 10 µM but remained non-toxic to TIGKs cells even at 50 µM (**Fig. 8B**). 350085 was able to inhibit the growth of HeLa cells from 5 µM upward and began showing toxicity to TIGKs at 20 µM, a four-fold higher concentration than what was observed on HeLa cells (**Fig. 8C**). Collectively, cell viability analysis indicate that cancerous cells are more sensitive to the LDH inhibitors than normal human cells and that our novel LDH inhibitors can inhibit the growth of HeLa cells at 5 – 10 µM.

**Figure 8.**
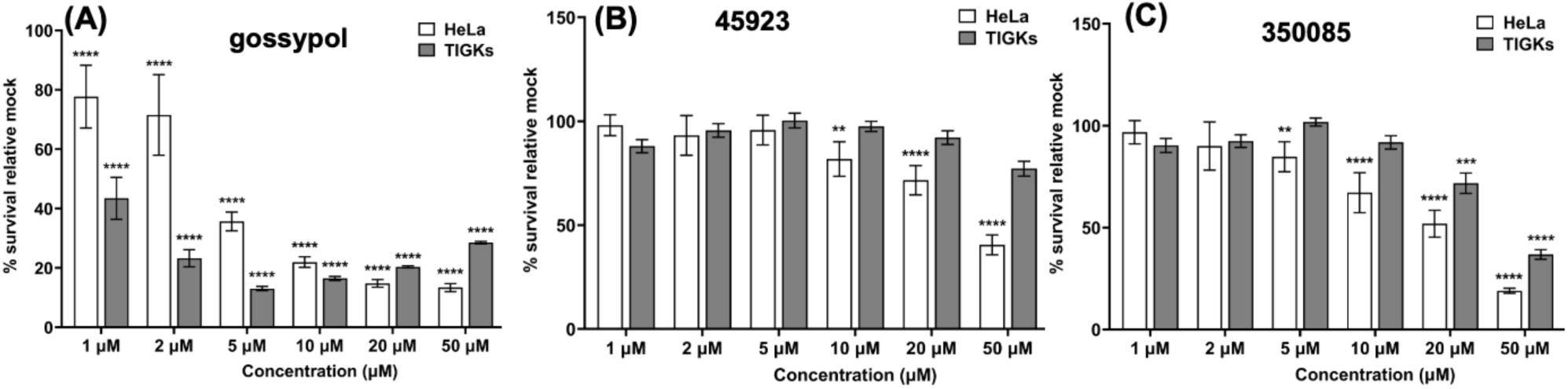
Impact of LDH inhibitors on the growth of HeLa and TIGKs cell lines. Growth analysis of an immortalized cervical cancer cell line, HeLa and telomerase-immortalized human gingival epithelial cell line (TIGKs) in the presence of **(A)** gossypol; **(B)** 45923; or **(C)** 350085. Cells were treated with the indicated concentration of inhibitors for 72 hours in a 96-well plate format and cell viability was determined using crystal violet assay as described (94). The growth inhibition study was performed in triplicate per concentration of inhibitors and in duplicate of technical and biological sets. Percentages of viable cells were normalized to mock treated wells and statistical analysis was performed using ANOVA. The results are expressed as means ± SEM from two to three biological replicates, significant difference (\**P* < 0.05; ** *P* < 0.01, *** *P* < 0.001, and **** *P* < 0.0001).

## Discussion

As a pathogen with limited genome capacity, *B. burgdorferi* relies heavily on its host to obtain essential nutrients such as fatty acids, amino acids, and nucleotides due to the absence of *de novo* biosynthesis pathways in its genome(21). Its restricted metabolic capacity leaves glycolysis as the sole source of ATP production(21, 52). LDH is the final enzyme in the glycolytic pathway, responsible for replenishing NADH by converting lactate, the end product of glycolysis, to pyruvate via the reduction of NAD^+^(30). As a tick-borne bacterium that alternates between an arthropod vector and mammalian host, *B. burgdorferi* must coordinate the uptake and utilization of different carbon sources, such as glucose and glycerol, to survive in diverse host environments. The genome of *B. burgdorferi* encodes a lactate permease (LctP, BB_0604), allowing the spirochetes to scavenge lactate from its external milieu(21). Lactate can be converted to pyruvate via BbLDH, altering the ratio of phosphoenolpyruvate to pyruvate, which signals the bacteria to utilize alternative carbohydrates for glycolysis. The expression of *lctP* is highly induced in the feeding tick vector, enabling a coordinated switch from glucose to glycerol as the primary carbon source(53). Ultimately, these carbon sources converge into the glycolysis pathway as substrates for ATP production(52). Beyond its role in carbohydrate metabolism, BbLDH can contribute to maintaining the intracellular pyruvate levels and protect the spirochetes against reactive oxygen species-mediated DNA damage and killing (27). Given its essential role in the energy supply chain and protective effects in maintaining redox balance and preventing oxidative killing, BbLDH represents a promising therapeutic target for Lyme disease. This potential can be extended to other tick- or louse-borne diseases caused by *Borrelia* species, as their LDHs shared high sequence identity (Extended Data Fig. 1).

Biochemical analyses reveal that BbLDH behaves similar to other LDHs, interconverting pyruvate (**Fig. 1A**) and lactate (**Fig. 1F**) through oxidation/reduction of NADH/NAD^+^. The addition of the allosteric activator FBP marginally reduced the *K_half_* of BbLDH for pyruvate (**Fig. 1A**). This finding is consistent with the crystal structure data, which show that FBP binding does not significantly alter the overall structure of BbLDH (**Fig. 2B**). Overall, the structures of the BbLDH tetramer with and without FBP (**Fig. 2A, B**) are very similar to the BsLDH tetramer (PDD: ILDB, Extended Data Fig. 4B), whereas the constituent dimers are also similar to the human LDH dimer (PDB: 8FW6, Extended Data Fig. 4A). LDH subunit conformations can be classified as either relaxed (R)- or taut (T)-state, with the allosteric regulator FBP favoring the R-state(36, 42). However, in BbLDH, FBP does not result in the tetramer-wide conformational conversion from the low-substrate affinity T-state to the high-substrate-affinity, R-state (**Fig. 2B**), largely because the FBP-minus structures are already in a primarily R-state conformation (Extended Data Fig. 5). The structural insensitivity to FPB may be a result of the high oxamate concentrations used in both crystallization conditions, which will favor the R-state conformation. Within the LDH tetramer, the four NADH/ oxamate do not deviate greatly from each other, with NADH and oxamate binding to several conserved residues, as seen in other LDH enzymes (**Fig. 2C**, **D**). However, close examination of the individual subunits reveals that their conformations are primarily distinguished only by a flexible loop adjacent to the NADH/oxamate binding site consisting of residues Gln85-Arg91 (Extended Data Fig. 4C). This loop is known as the activity site A-loop as it sits above the active site pocket and has been shown to undergo conformational changes during substrate processing(54). The two FBP-bound BbLDH dimers within the tetramer are practically identical, except for the relative position of the A-loop (Extended Data Fig. 2D). Hence in BbLDH, the A-loop is conformationally variable regardless of the presence of FBP. Previously, it has been shown that the A-loop adopts an open conformation in the T-state and a close conformation in the R-state in *Bifidobacterium longum* subsp. *Longum* LDH (BlLDH)(42), although these two conformations have also been observed in non-allosterically controlled LDH enzymes(54). Comparing the A-loop position between the apo-FBP and FBP-bound LDH structures and the R- and T-state of BlLDH subunits supports the assertion that BbLDH adopts an R-state conformation independent of FBP, at least in the crystallized state (Extended Data Fig. 5). Although BbLDH crystallized as a tetramer similar to other bacterial LDH enzymes, solution-state measurements revealed that BbLDH is a dimer in solution, regardless of the presence of oxamate, NADH, FBP, or any combination of the three (Extended Data Fig. 6). Thus, either BbLDH functions primarily as a dimer, like HsLDH, or it tetramerizes under higher local cellular concentrations or in the presence of additional factors.

Bacterial LDHs have been implicated in the virulence and pathogenesis of several bacterial species(55–58). For instance, in *Enterococcus faecalis*, the ability of the bacteria to resist various environmental stresses relies on the role of LDH in maintaining redox balance. Deficiency in LDH leads to increased susceptibility to various stressors and diminished capacity to colonize murine hosts(57). In *Streptococcus pneumoniae*, the loss of LDH activity renders the pathogen avirulent, likely due to inefficiencies in metabolism and virulence gene expression(55). We were unable to achieve a complete knockout of BbLDH in *B. burgdorferi* and using IPTG-inducible strain we confirmed that BbLDH is essential for the growth of the spirochetes (**Fig. 3D**). This phenotype is distinct from those observed in the LDH mutants of other bacteria, which typically carry more than one copy of *ldh* gene in their genomes(55) and can utilize alternative catabolic pathways, such as the Tricarboxylic Acid Cycle (TCA) or Pentose Phosphate Pathway (PPP),(59, 60) to synthesize ATP. In contrast, *B. burgdorferi* encodes only a single copy of *ldh* and lacks the genes required for alternative metabolic pathways like the TCA and PPP(21). This limitation underscores why the deletion of BbLDH is lethal to the spirochete, as glycolysis becomes the sole energy production pathway (**Fig. 3**).

A follow-up mouse infection study indicated that BbLDH is important for the pathogenicity of *B. burgdorferi*, as the deletion of BbLDH significantly attenuated the ability of the spirochetes to cause systemic infection and elicit a strong host immune reaction (**Fig. 4**). Even though BbLDH is essential for growth in vitro, several factors may contribute to the ability of the *ldh* mutant spirochetes disseminating to some extent *in vivo.* Prolonged infection could have induced spontaneous mutations that suppressed *lacI* expression from the shuttle vector, allowing leaky expression of BbLDH. Furthermore, *B. burgdorferi* may be able to obtain host factors that partially compensate for the loss of BbLDH. Nonetheless, there is a striking, ∼5-fold higher level of spirochete burden when the mice drink the IPTG inducer. This result is quite remarkable given that we were unable to control the amount of IPTG consumed by each mouse, as the drinking habits varied among individuals. Consequently, the expression level of BbLDH induced *in vivo* can differ between mice. Second, the stability of IPTG in the drinking water over three-week infection study may have been compromised, leading to reduced induction efficiency in BbLDH expression. Lastly, IPTG has to survive metabolic processing in the mouse, traverse the vascular system and permeate tissues to reach the spirochetes and induce gene expression within them. Thus, the ∼ 5-fold higher level of spirochetes burden and a detectable host immune reactions compared to those that received regular drinking water, is strong evidence for the importance of BbLDH to spirochete viability *in vivo* (**Fig. 4A**).

From the natural compound library screening, four inhibitors with anti-LDH activities were identified and all four compounds are structurally distinct from the well-established LDH inhibitor, gossypol (**Fig. 5A**). Two of the four compounds, 45923 (Methoxsalen) and 350085 (Medicarpin), have been extensively used in animal models for studies unrelated to LDH. Methoxsalen is used to treat psoriasis in combination with ultraviolet A (UVA) light therapy (Puvatherapy)(61–63) and for vitiligo(64). In these applications, photoactivated methoxsalen alkylates DNA leading to formation of DNA adducts(65, 66). Without photoactivation, methoxsalen was reported to induce mutation in bacteria but the data was inconclusive(67). Medicarpin on the other hand is a bioactive pterocarpan(68), a group of isoflavonoids with diverse biological functions including anti-inflammatory to antimicrobial properties(69–75). Anti-osteoporotic and bone protection benefits have been reported with medicarpin in animal models(73, 76). Cytotoxic effects have also been observed upon treatment of cancer cells with medicarpin(77, 78) along with antifungal(79) and antibacterial properties(80). In this study, methoxsalen was shown to have anti-LDH activity without activation using UVA radiation (**Fig. 5D**) as well as having anti-proliferation effects on *B. burgdorferi* (**Fig. 6C**) and cancerous cells, but not normal cells (**Fig. 8B**). The inhibitory effect of medicarpin on the growth of *B. burgdorferi* was not as potent as methoxsalen (**Fig. 6E**). This observation could be impacted by the solubility of each compound in BSK-II growth media and variation in the permeability of each compound to *B. burgdorferi*. Future studies will be directed towards increasing the potency of these compounds by improving their solubilities as well as their uptake by the bacteria. In addition to inhibition of *B. burgdorferi* growth, with medicarpin at 5 µM has cytotoxic effects on HeLa cells, yet was well-tolerated by TIGKs cells at up to four times of this concentration (**Fig. 8C**). To the best of our knowledge, this is the first report of methoxsalen and medicarpin exhibiting anti-LDH activities.

We were unable to determine the crystal structures of BbLDH bound to gossypol, 14975, 45923, 114344 or 350085. Although these inhibitors clearly interact with the protein in solution, their binding may be incompatible with the tetramer of BbLDH that preferentially forms in the crystal. Thus, the crystallization of BbLDH may occlude the inhibitor depending on where and how the compounds interact with LDH. To circumvent this issue, we docked each inhibitor to BbLDH *in silico*. Lineweaver–Burk plots of each inhibitor against pyruvate suggested that gossypol is an uncompetitive inhibitor while 14975, 45923, 114344 and 350085 are competitive inhibitors (Extended Data Fig. 7B, C). Therefore, gossypol was docked to an LDH model containing NADH and oxamate, while 14975, 45923, 114344 and 350085 were docked to an apo LDH model. Recently, the crystal structure of HsLDH in complex with gossypol has been determined to a resolution of 3.5Å. This structure shows that gossypol binds to the A-loop of HsLDH and interacts with residues Glu103, Asn107, Gln110 and Arg111. Our docking analysis of gossypol to BbLDH revealed a similar binding mode and set of interacting residues, thereby validating this approach for predicting the general binding configurations of 14975, 45923, 114344 and 350085 (**Fig. 7A, B**). In BbLDH, gossypol binds to an anionic pocket formed by the A-loop (**Fig. 7A**) and hydrogen bonds to Lys86 and Asp96 (**Fig. 7B**). The docking pose of gossypol indicates that its binding is compatible with the closed A-loop conformation of the R-state but not the open configuration of the T-state (Extended Data Fig. 8). This observation lends support to the uncompetitive inhibition observed for gossypol because the cofactor NADH and substrate lactate/pyruvate drive A-loop closure and conversion to the R-state.

We next examined the top docking pose of each of the candidate inhibitors identified in this study. All four inhibitors bind to the NADH/lactate binding pocket with docking scores ranging from -4.124 to -4.801 (Extended Data Figs. 9-10). Each inhibitor interacts with a slightly different set of residues, although there are some commonalities. For example, all three of 14975, 45923, and 114344 form hydrogen bonds with a span of glycine residues: Gly13, Gly14 and Gly16. The most potent inhibitor, 45923, makes only one hydrogen bonding contact with Gly14 in BbLDH, whereas both the least potent inhibitor and the second most potent inhibitor form four hydrogen bonding contacts with this region. Other factors that may also contribute to inhibition by these compounds, include the burying of hydrophobic surface area and interactions with residues Val38, Leu83, Thr80, and the Gly11-Ala12-Gly-13-Gly14-Val15-Gly16 peptide motif. Ultimately, these structural models enabled by the BbLDH crystal structures provide testable hypotheses of binding modes that can be further improved upon by future compound derivatization.

In summary, our findings establish BbLDH as a functional lactate dehydrogenase enzyme essential for the survival and pathogenesis of *B. burgdorferi*. We have identified four novel small compounds with anti-LDH activities, which provide a solid foundation for further investigation into the feasibility of targeting BbLDH to curb the spread of LD. With the determined crystal structure of BbLDH and the potential that it provides for rational design guided by computation, future work will focus on refining the specificity and potency of these compounds as metabolic inhibitors against *B. burgdorferi*.

## Methods

### Bacterial strains, tissue culture cell lines and growth conditions

Infectious clone B31 A3-68 (wild type), a derivative strain from the *B. burgdorferi* sensu stricto B31-A3, was used in this study(46). Cells were grown in Barbour-Stoenner-Kelly (BSK-II) medium as previously described (81) in appropriate antibiotic(s) for selective pressure as needed: streptomycin (50 µg/ml), kanamycin (300 µg/ml), and/or gentamicin (40 µg/ml). Cells were maintained at 34°C, pH 7.4 in the presence of 3.4% CO_2_.

*Escherichia coli* DH5α strain (New England Biolabs, Ipswich, MA) was used for DNA cloning and Rosetta-gami (DE3) (Novagen, San Diego, CA) was used for recombinant protein expression. *E. coli* strains were grown in lysogeny broth (LB) supplemented with appropriate concentrations of antibiotics for selective pressure as needed: ampicillin (100 µg/ml), chloramphenicol (34 µg/ml), streptomycin (50 µg/ml), tetracycline (12.5 µg/ml), and spectinomycin (50 µg/ml).

Immortalized cervical carcinoma cells (HeLa) and Telomerase Immortalized Gingival Keratinocytes (TIGKs) were purchased from ATCC. HeLa cells were cultured and maintained in DMEM (Dulbecco’s Modified Eagles Medium) with high glucose and supplemented with 10% FBS and 100µg/ml of penicillin and streptomycin. TIGKs cells were cultured in serum-free keratinocyte growth medium (KGM; Lifeline Cell Technology, MD) as described previously(51). Cells were incubated at 37°C, 5% CO_2_ incubator.

### Construction of pJSB-BB_0087FLAG and BB_0087::kan for conditional knockout in *B. burgdorferi*

The inducible shuttle vector pJSB275 was a kind gift from Dr. Blevins(47). In order to clone *bb_0087* into the inducible vector, we first modified the vector and expanded the multiple cloning sites beyond NdeI and HindIII to include XhoI-BamHI-NruI-SacII. The open reading frame (*orf*) of *bb_0087* with a C-terminal FLAG was PCR amplified from wild-type B31 A3-68 using the primer pair P_1_/P_2_ and cloned into the modified pJSB275 at the engineered XhoI and SacII sites, yielding pJSB-BB_0087FLAG (**Fig. 3A**). To inactivate the endogenous *bb_0087* gene in *B. burgdorferi* via allelic exchange mutagenesis, the upstream and downstream flanking regions of *bb_0087* were PCR amplified using the primer pairs P_3_/P_4_ and P_5_/P_6_, respectively. The kanamycin resistance cassette (*kan*) was PCR amplified using primer pair P_7_/P_8_. The three amplicons were then PCR ligated using primer pair P_3_/P_6_ and cloned into pGEM-T easy vector, yielding BB_0087::kan (**Fig. 3B**). All primers used in this study were synthesized from IDT (Integrated DNA Technologies, Coralville, IA) and listed in Extended Data Table 1.

**Table 1.** Data collection and refinement statistics (molecular replacement)

|  | BbLDH + FBP (9DQ7) | BbLDH – FBP (9DQ9) |
| --- | --- | --- |
| **Data collection** |  |  |
| Resolution range (Å) | 48.98-2.11 (2.18-2.11) | 47.47-2.10 (2.18-2.10) |
| Space group | P2<sub>1</sub>2<sub>1</sub>2<sub>1</sub> | P2<sub>1</sub>2<sub>1</sub>2<sub>1</sub> |
| Unit cell dimensions<br>a,b,c<br>$\alpha, \beta, \gamma$ | 81.0, 91.2, 166.6<br>90, 90, 90 | 80.5, 90.4, 167.4<br>90, 90, 90 |
| Total reflections | 440210 | 536124 |
| Unique reflections | 70826 (6558) | 71544 (7019) |
| Multiplicity | 6.2 (5.7) | 3.4 (3.1) |
| Completeness (%) | 98.7 (92.2) | 97.9 (85.4) |
| Mean I/s(I) | 12.7 | 3.7 |
| Wilson B-factor | 28.74 | 42.83 |
| R-merge/meas/pim | 0.237/0.259/0.103 | 0.361/0.393/0.154 |
| CC1/2 | 0.97 | 0.80 |
| **Refinement** |  |  |
| Reflections use for refinement | 70764 (6513) | 70464 (6070) |
| Reflections used for R-free | 2000 (185) | 1882 (162) |
| R-work | 0.1782 (0.239) | 0.2018 (0.349) |
| R-free | 0.2211 (0.288) | 0.2130 (0.360) |
| Number of non-hydrogen atoms | 10389 | 10133 |
| Macromolecules | 9737 | 9748 |
| Ligands | 378 | 321 |
| Solvent | 410 | 180 |
| Protein Residues | 1257 | 1258 |
| RMS (bonds) | 0.007 | 0.005 |
| RMS (angles) | 0.86 | 0.72 |
| Ramachandran favored (%) | 95.9 | 94.0 |
| Ramachandran allowed (%) | 3.9 | 5.6 |
| Ramachandran outliers (%) | 0.16 | 0.40 |
| Rotamer Outliers (%) | 1.8 | 1.7 |
| Clashscore | 4.5 | 4.9 |
| Average B-factor | 35.3 | 49.5 |
| Macromolecules | 35.2 | 49.6 |
| Ligands | 34.0 | 49.8 |
| Solvent | 38.1 | 46.1 |
| - Statistics for the highest-resolution shell are shown in parentheses.<br>- Both data sets were collected at the Cornell High Energy Synchrotron Source (CHESS)<br>- Data sets were collected at 77K at a wavelength of 0.9686 Å |  |  |

### Enzyme kinetics and inhibition assays

To establish the LDH activity of BbLDH, we first used a commercial Lactate Dehydrogenase Activity Assay Kit (Sigma Aldrich, St. Louis, MO) to measure the ability of BbLDH to reduce NAD to NADH following the manufacturer’s instruction. Commercial LDH inhibitor gossypol (catalog # G8761) was dissolved in DMSO at a stock concentration of 100 mM (Sigma Aldrich). 1 µg/µl of recombinant wild-type or H178A BbLDH stocks were diluted 1:20 in the LDH assay buffer to a final working volume of 400 µl per well. 100 µM of gossypol or DMSO were added to selected wild-type BbLDH wells to determine the inhibitory effect of gossypol on BbLDH activity. Reactions were set up in triplicate wells along with NADH standards. The NADH activity was calculated using the formulation provided by the manufacturer.

The enzyme kinetics of BbLDH, with substrates pyruvate or lactate, was monitored by the reduction or oxidation of NADH at OD_340_ as described(82) using a Varioskan LUX multimode microplate reader (Thermo Fisher Scientific, Rockford, IL) with minor modification. Briefly, the conversion of pyruvate to lactate activity was performed at 37°C in 50 mM Tris-HCl buffer pH 7.0 containing 100 ng of BbLDH, 0.5 mM NADH, 2.4 mM pyruvate and with/without 3 mM fructose 1,6-bisphosphate (FBP). The conversion of lactate to pyruvate activity was measured at 37°C in sodium bicarbonate buffer pH 7.0 containing 100 ng of BbLDH, 1 mM NAD^+^, 0 – 300 mM sodium L-lactate, with/without 3 mM FBP. The kinetic parameters of BbLDH were determined using 0 – 6.0 mM pyruvate, 0 – 1.0 mM NADH, 0 – 2.0 mM NAD^+^, 50 mM Tris-HCl buffer pH 5.0 - 10.0, or reaction temperature of 25°C, 30°C, and 37°C. All reactions were performed in 96-well plate with a final volume of 200 µl per well.

The inhibitory effect of gossypol and the four lead inhibitors of BbLDH was performed using 0 – 100 µM of inhibitors or equal volume of DMSO vehicle in 50 mM Tris-HCl buffer pH 7.0 containing 100 ng of BbLDH, 0.5 mM NADH, 2.4 mM pyruvate and 3 mM FBP at 37°C. All experiments were repeated at least twice. The amount of NADH produced or consumed were extrapolated from the NADH standard curve and plotted in GraphPad Prism 10 (GraphPad Software, La Jolla, CA, USA) using Allosteric sigmoidal mode to obtain the *V*_max_ and *K*_half_ values. Lineweaver-Burk plots were generated using the mean reciprocal values of substrates and velocities obtained from the inhibition studies.

### Crystallization and structure determination of BbLDH

BbLDH was prepared in 20 mM Tris pH 8.0, 100 mM NaCl to a final concentration of 10 mg/mL and supplemented with 1 mM NADH, 1 mM oxamate, and 1 mM FBP, the latter only for the FBP-bound state. To crystallize BbLDH+FBP and BbLDH – FBP, 1 μL of 10 mg/mL BbLDH was mixed with 1 μL 100 mM HEPES pH 7.4-7.6, 100 mM calcium acetate, and 40% (m/v) PEG400 in hanging well plates. Crystals appeared after 5 days at room temperature. Data sets were collected at the Cornell High Energy Synchrotron Source (CHESS) beam-line ID7B2 at 77K with an X-ray wavelength of 0.9686 Å. Data sets were integrated and scaled in HKL2000 and SCALEPACK. The structure was determined by molecular replacement using a preliminary lower-resolution BbLDH homology model as a probe. Manual adjustments were made with COOT and refinement was carried out with Phenix Refine. The final BbLDH+FBP and BbLDH-FBP structures had a Rwork/Rfree (%/%) of 0.178/0.221 and 0.202/0.213, respectively. The Ramachandran and rotamer outliers (%) for each structure are as follows (structure-Ram. outliers/rotamer outliers): BbLDH+FBP-0.16/1.80, BbLDH-FBP-0.40/1.70. Statistics for data processing and refinement are reported in Table 1. Figures were prepared in COOT and PyMol. BbLDH+FBP and BbLDH-FBP structures were deposited to the PDB with the accession ID 9DQ7 and 9DQ9, respectively.

### Docking analysis protocol

To generate BbLDH models for docking analysis, the crystal structure of BbLDH without FBP (PDB ID: 9DQ7) was used as a starting model. The TS LDH dimer was imported into Maestro(83) and the solvent water molecules and calcium ions were removed. For the apo LDH model, NADH and oxamate were also removed. Each protein was prepared using an identical workflow. First, the models were prepared using protein preparation wizard(84, 85). A pH of 7.0 was used based on the experimental conditions of our inhibition assays. Prepared model energy minimization was performed using Macromodel (86) minimization using the following parameters: OPLS4 force-field (87), implicit solvation with a constant dielectric of 78, charges were calculated from the force-field with normal cutoffs. PRCG minimization method, 100,000 maximum iterations and models were allowed to converge on a gradient with a threshold of 1×10^-4^. Ligands were imported and prepared using LigPrep(88) software at a pH of 7. Docking grids (20 x 20 x 20 Å^3^) were generated in Glide(89) centered on residue Asp96, encompassing the NADH, FBP and lactate binding sites in both models. Docking poses were generated using Glide Dock using default parameters with modifications. For each ligand, ligands were dock using standard-precision. Ligand-protein hydrogen bonds were rewarded with at least 2000 poses considered per ligand. All poses were subject to post-docking energy minimization using a OPLS4 force field(87). A maximum of 2,000 poses per ligand were considered and the top pose output for analysis.

### Measuring the growth rates of *B. burgdorferi*

To measure the growth of the WT B31 A3-68 and 87^mut^, 87^mut^ was first cultivated in the presence of 1 mM IPTG until stationary-phase along with the WT. Cells were harvested via centrifugation and washed with PBS to remove all traces of IPTG. 1 x 10^5^ cells/ml of WT or 87^mut^ were inoculated into 10 ml of fresh BSK-II and incubated at 34°C with or without 1 mM IPTG. Bacterial cells were enumerated daily until the cells entered the stationary growth phase (∼1 x 10^8^ cells/ml) using a Petroff-Hausser counting chamber as previously described(90). At the end of the growth study, cultures were harvested for immunoblot analysis to detect for the expression of BbLDH. Growth analyses were repeated in triplicates and the results were expressed as the means ± standard errors of the means (SEM). To determine the growth inhibition of LDH inhibitors on *B. burgdorferi*, 10^5^ cells/ml of stationary WT cultures were inoculated into fresh 5 ml BSK-II media containing the specified concentration of inhibitors. Equal volume of DMSO were included as negative controls to rule out the inhibitory effect of DMSO on the growth of *B. burgdorferi*. Cells were maintained at 34°C with a 3.4% CO_2_ incubator and cell numbers were enumerated every two days until the cells reach the stationary growth phase using Petroff-Hausser counting chamber.

### Electrophoresis and immunoblotting analyses

For the detection of recombinant protein inductions and purification, equal amounts of *E. coli* whole cell lysates or recombinant proteins were separated on 10% SDS-PAGE gel followed by Coomassie blue staining (0.05% Coomassie brilliant blue R250, 25% isopropanol, 10% acetic acid). To detect the expression of inducible BbLDH, 10 to 20 μg of *B. burgdorferi* whole cell lysates were separated on 10% SDS-PAGE gel and transferred to PVDF membrane (Bio-Rad Laboratories, Hercules, CA). The immunoblots were probed with polyclonal antibodies against *B. burgdorferi* BbLDH. DnaK was used as an internal control, as previously described (81). Membranes were developed using horseradish peroxidase secondary antibody with an ECL luminol assay or with fluorescently labelled secondary antibodies. Signals were imaged using the ChemiDoc MP Imaging System and analyzed using Image Lab software (Bio-Rad Laboratories). For mouse seroconversion test, wild-type *B. burgdorferi* lysates were analyzed on a 12% SDS-PAGE and transferred to PVDF membrane as described above. After membrane blocking using 5% non-fat milk, the PVDF membrane containing WT lysate was assembled onto the Mini-PROTEAN II Multiscreen Apparatus (Bio-Rad Laboratories). 600 µl of 1: 100 diluted mouse sera in 2% non-fat milk were loaded into separate channel and incubated at 4°C overnight with gentle shaking. Sera from two mock-infected mice were included as negative controls. Following overnight incubation, immunoblots were performed and developed as described.

### Mouse infection studies

BALB/c mice at 6-8 weeks of age (Jackson Laboratory, Bar Harbor, MN) were used in the needle infection study. All animal experimentation was conducted following the NIH guidelines for housing and care of laboratory animals and was performed in accordance with the Virginia Commonwealth University institutional regulations after review and approval by the Institutional Animal Care and Use Committees. The animal studies were carried out as previously described(91, 92) with minor modifications. To prepare *B. burgdorferi* cells for animal study, WT and 87^mut^ cells were cultured in the presence of 1 mM IPTG to mid log phase (∼ 10^6^ cells/ml). Cells were harvested and washed three times in PBS before being reinoculated into fresh BSK-II media without IPTG and cultured for another three days to ensure depletion of all IPTG from the cells.

Mice were given a single subcutaneous injection of 10^3^ spirochetes in 100 µl volume. Mice injected with 87^mut^ were separated into two groups, one group received normal drinking water while the other received IPTG-supplemented water (sterilized 2% sucrose solution containing 80 mM IPTG) as described previously(93). Mice were sacrificed 3 weeks post-infection. Ear tissues and whole blood were collected for qRT-PCR analysis and seroconversion test. Tissues from the ears were harvested to measure the bacterial burdens using qRT-PCR, as previously described(81). Briefly, total RNAs from the mouse tissues were isolated using TRIzol reagent (Invitrogen, Carlsbad, CA) and contaminating genomic DNA was removed using DNase (Takara Bio USA, Mountain View, CA). The DNase-treated RNAs were reverse transcribed to cDNA using SuperScript IV VILO Master Mix (Thermo Fisher Scientific) according to the manufacturer’s instructions. Quantitative PCR (qPCR) was performed using Fast SYBR Green Master Mix (Applied Biosystems, Foster City, CA) with primers targeting the *flaB* and mouse *β-actin* as described previously(91). The spirochete burdens within the infected ear tissues were expressed as the copy number of *flaB* transcript per 10^5^ of mouse *β-actin* transcript.

### Small compound library screening

To identify potential novel LDH inhibitors against BbLDH, the Natural Products Set IV library containing 419 natural compounds selected from the DTP Open Repository collection was obtained from the National Cancer Institute (NCI). All compounds were dissolved in DMSO to a working stock solution of 10 mM. Primary screening was performed using a label free high throughput screening (HTS) protocol(49) in 384-well plate format. Buffer B (20 mM Tris pH 7.4, 0.005% Triton X-100, 0.005% BGG, 2 mM DTT) containing 10 nM BbLDH, 75 µM pyruvate, and 50 µM NADH was dispensed into each well containing pre-dispensed inhibitor to a final concentration of 10 µM in 50 µl working volume per well. Fluorescence was measured using an excitation wavelength of 340 nm and an emission wavelength of 482 nm. Readings were recorded every second over a total duration of 5 minutes. Gossypol and DMSO were included as positive and negative controls, respectively. Fluorescent signals were plotted over time for each compound and analyzed using linear regression to determine the slope of reduction compared to DMSO and gossypol. Twenty three compounds with a reduced slope in comparison to DMSO were selected for a second round of screening.

For the secondary screening, Lactate Dehydrogenase Activity Assay Kit (Sigma Aldrich) was used to determine the inhibitory effect of selected small compounds on BbLDH activity. Reactions were performed following the manufacturer’s protocol using 10 and 25 µM of small compounds and 10 nM BbLDH per well. Gossypol and DMSO were included as positive and negative controls, respectively. Compounds that reduced the LDH activity of BbLDH compared to DMSO treatment well were selected for further analysis. Of the 23 compounds, 14975, 45923, 114344, and 350085 showed the highest inhibition on BbLDH activity.

### LDH inhibitor and cell viability test

To determine the toxicity of LDH inhibitors on eukaryotic cells, HeLa and TIGKs cells were seeded onto 96-well plate at 10^4^ cells per well the day before drug treatment. Cell culture media was removed the next day and replaced with culture media containing the specified concentration of DMSO, gossypol, 45923, or 350085. Mock-treated wells were included for cell viability normalization purposes. Cells were incubated with inhibitors for 72 hours. To determine the cell viability, crystal violet assay was used as described(94). Briefly, cell culture media were decanted from the 96-well plate and washed once with water. 50 µl of 0.5% crystal violet staining solution (0.5 g crystal violet powder, 80 ml ddH_2_O, 20 ml methanol) and incubated for 20 min at room temperature (RT) with gentle shaking. Plates were washed 4-5 times by soaking in ddH2O followed by gentle tapping to remove any remaining liquid. Plates were air dried at RT. 200 µl of methanol was added to each well and plates were incubated for 20 min at RT with gentle shaking followed by optical density measurement at 570 nm (OD_570_) using a Varioskan LUX multimode microplate reader (Thermo Fisher Scientific). The OD_570_ of mock cells were set to 100% and the percentage of viable cells treated with inhibitors were determined by comparing the average OD_570_ values of treated wells to mock wells. Experiment was performed in triplicate wells with two technical replicates and two biological replicates. Two way ANOVA was used for statistical analysis with significant values of *P* < 0.05.

## Acknowledgments

We thank Dr. Blevins for providing pJSB275 vector. This work was supported by funding from the National Institutes of Allergy and Infectious Diseases (AI078958 to C. Li; AI148844 to B. Crane and C. Li), National Institutes of Health (NIH). This work is based on research conducted at the Center for High-Energy X-ray Sciences (CHEXS), which is supported by the National Science Foundation (BIO, ENG and MPS Directorates) under award DMR-1829070, and the Macromolecular Diffraction at CHESS (MacCHESS) facility, which is supported by award 1-P30-GM124166-01A1 from the National Institute of General Medical Sciences, National Institutes of Health, and by New York State’s Empire State Development Corporation (NYSTAR).

## Data availability

The data and detailed protocols that support the findings of this study are available without restriction from the corresponding authors. Source data are provided with this paper.

## Author contributions

C.L. and B.R.C. secured the funding, designed the project, supervised the research, coordinated the collaboration, and wrote the manuscript. C.W.S, M.J.L., and K.Z. designed, conducted, and analyzed the experimental work and are responsible for data collection and manuscript preparation. D.N., S.E., M.J.L and B.R.C. conducted X-ray crystallography study and data collection.

## Additional Information

Extended Data (Methods, 10 figures, and 4 tables).

## Supplementary Data

### Methods

#### Recombinant Protein Expression & Purification

The *orf* (*bb_*0087) for BbLDH was PCR amplified from wild-type B31 A3-68 using primer pair P_1_/P_9_ with the XhoI and SacII restriction sites engineered to the 5’ and 3’ end, respectively. The resultant amplicon was cloned into pGEM-Teasy vector and excised with XhoI and SacII for ligation into a modified pET100/D-TOPO expression vector with an N-terminal 10 x His 1 x FLAG tag, yielding pET-BB_0087. The resulting pET-BB_0087 plasmid was transformed into Rosetta-gami 2 (DE3) *E. coli* strain (Novagen). For protein induction, 20 ml of overnight starter culture was inoculated into 1 L of LB broth and grown to OD_600_ of 0.5 at 37°C with agitation. The expression of recombinant BbLDH was induced using 1 mM isopropyl-β-D-thiogalactoside (IPTG) at 37°C for four hours and then purified at 4°C using HisTrap HP columns following the manufacturer’s instruction. A second purification step using Size Exclusion Chromatography (SEC) was added to purify the eluted protein further and dialyzed in 10 mM Tris buffer, pH 8.0 at 4°C overnight. The purified recombinant protein was aliquoted into smaller volumes and stored at -80°C for antibody production, enzymatic analysis, and crystallization. To study the significance of key residues in the function of BbLDH, pET-BB_0087 constructed above was used as template for all the site directed mutagenesis study. pET-BB_0087H178A and pET-BB_0087T146A R154A were constructed using primer pairs P_10_/P_11_ and P_12_/P_13_, respectively, to establish the role of BB_0087 as a LDH. pET-BB_0087H171A was constructed using primer pair P_14_/P_15_ to determine if BbLDH is allosterically regulated.

#### Production of BbLDH polyclonal antibodies

5 mg of purified recombinant wild-type BbLDH protein from above was used to generate polyclonal antibodies in rabbits on a fee-for-service basis in General Bioscience Corporation (Brisbane, CA), following a standard immunization procedure.

#### SEC-MALS of BbLDH with and without cofactors

BbLDH samples were prepared to a final concentration of 10 μM in MALS buffer (20 mM Tris pH 7.5, 150 mM NaCl) supplemented with 10 mM NADH, oxamate and/or NADH. Prior to injection, 10 mM β-mercaptoethanol was added to each sample and centrifuged at 13,000 rpm for 5 minutes at 4°C. Each sample (100 mL) was injected onto a S200 (10/30, GE Lifesciences) that had been pre-equilibrated with MALS buffer for >3 CVs. The gel filtration column was coupled to a static 18-angle light scattering detector (DAWN HELEOS-II) and a refractive index detector (Optilab T-rEX) (Wyatt Technology, Goleta, CA). Data were collected every second at a flow rate of 0.7 mL/min. Data analysis was carried out using ASTRA VI, yielding the molar mass and polydispersity of the sample. Monomeric BSA was used to standardize the light scattering detector.

**Table S1.**
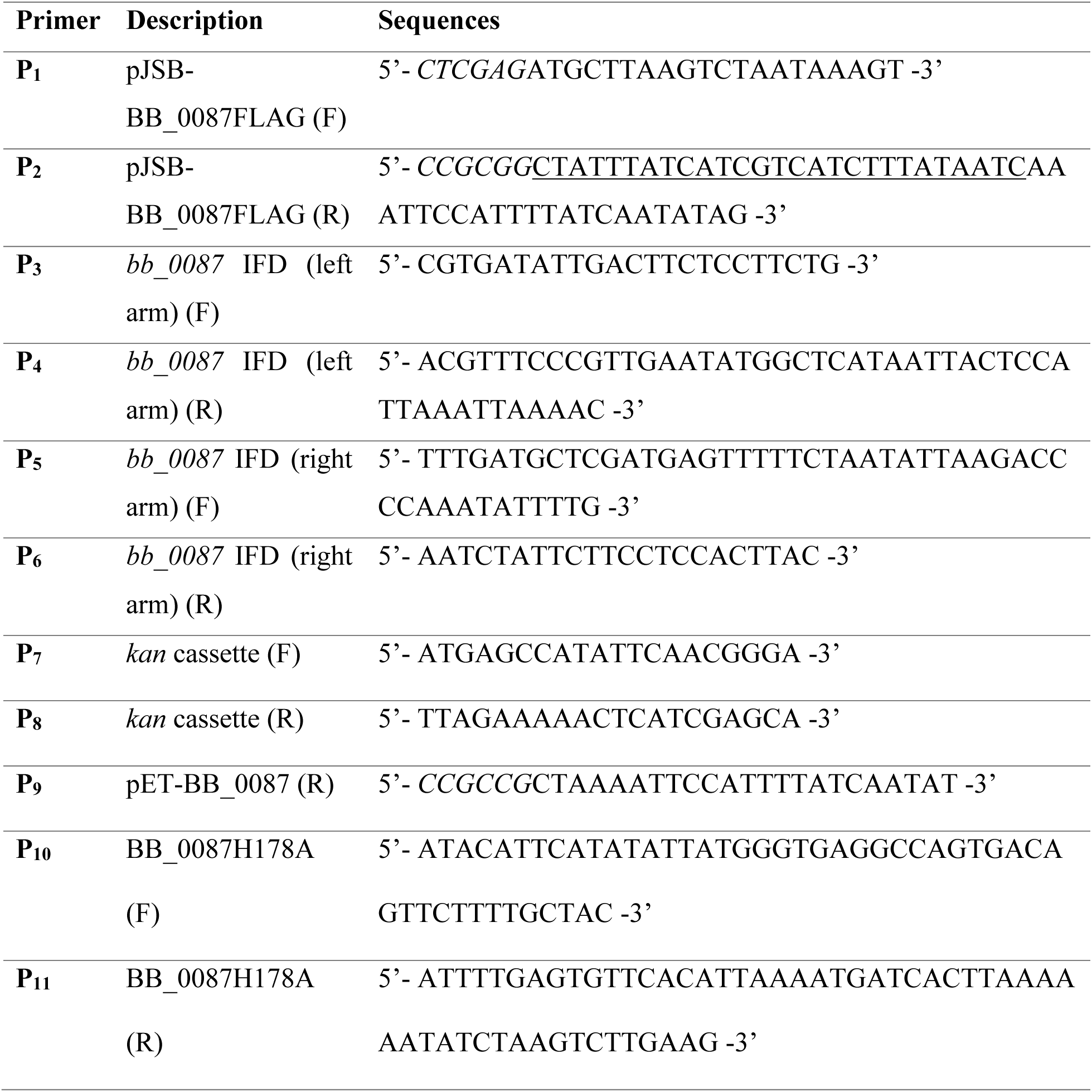
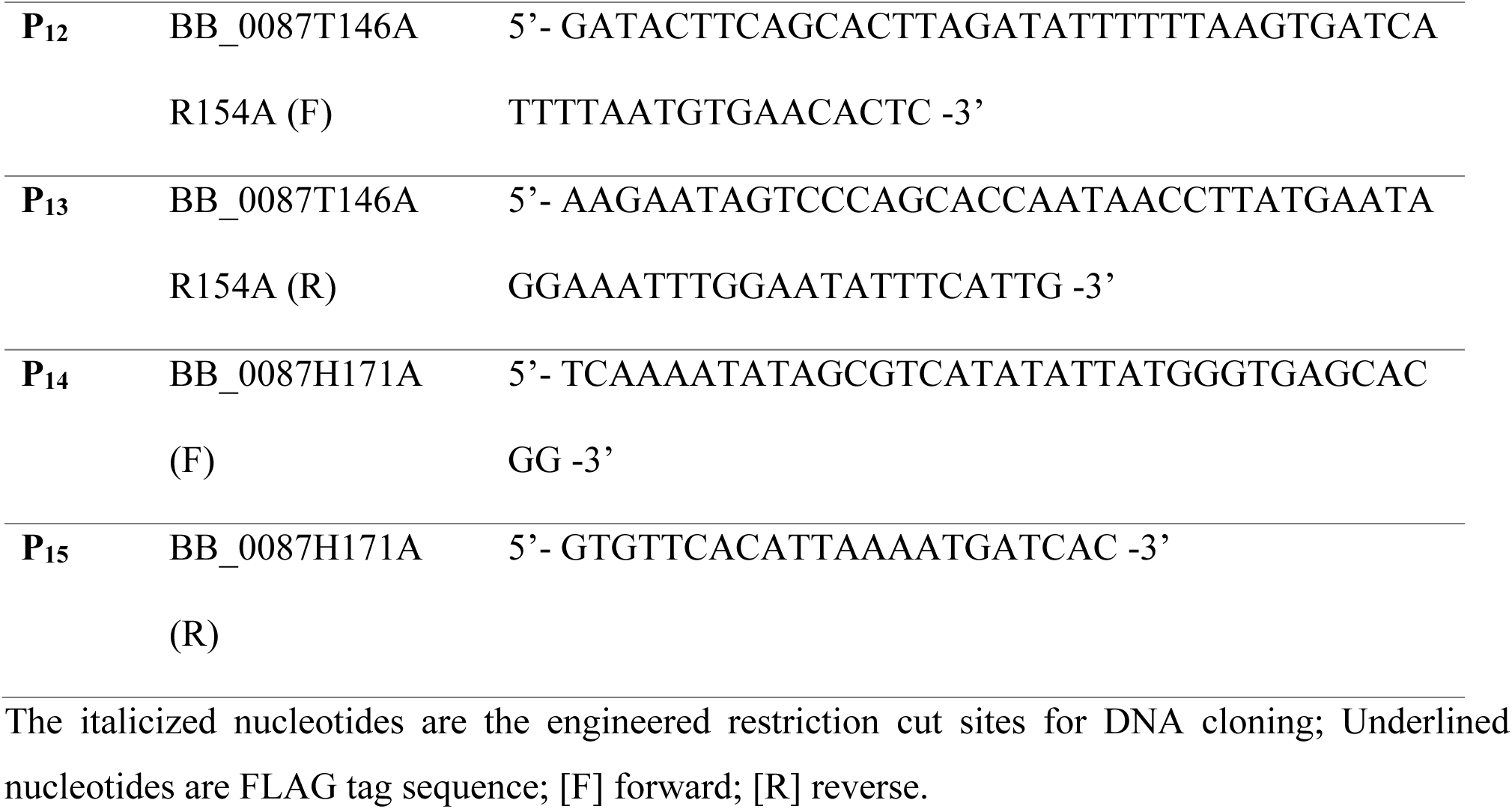
Oligonucleotide primers used in this study.

**Table S2:** Interactions between LDH (chain A) and NADH.

| Index | Interaction | Residue | Distance (Å) |
| --- | --- | --- | --- |
| 1 | hydrophobic | L150 | 3.93 |
| 2 | hydrophobic | I235 | 3.74 |
| 3 | H-bond | A12 | 4.0 |
| 4 | H-bond | G14 | 3.3<br>3.1 |
| 5 | H-bond | V15 | 3.1 |
| 6 | H-bond | D37 | 2.6 |
| 7 | H-bond | D37 | 2.7 |
| 8 | H-bond | V38 | 3.5 |
| 9 | H-bond | Y68 | 4.0 |
| 10 | H-bond | G82 | 3.4 |
| 11 | H-bond | N84 | 3.1 |
| 12 | H-bond | N98 | 4.1 |
| 13 | H-bond | A121 | 3.1 |
| 14 | H-bond | N123 | 3.1<br>3.2 |
| 16 | H-bond | G13 | 3.7 |
| 17 | H-bond | T146 | 2.9 |
| 18 | H-bond | water | 2.9<br>2.8 x 4<br>2.7 |

**Table S3:**
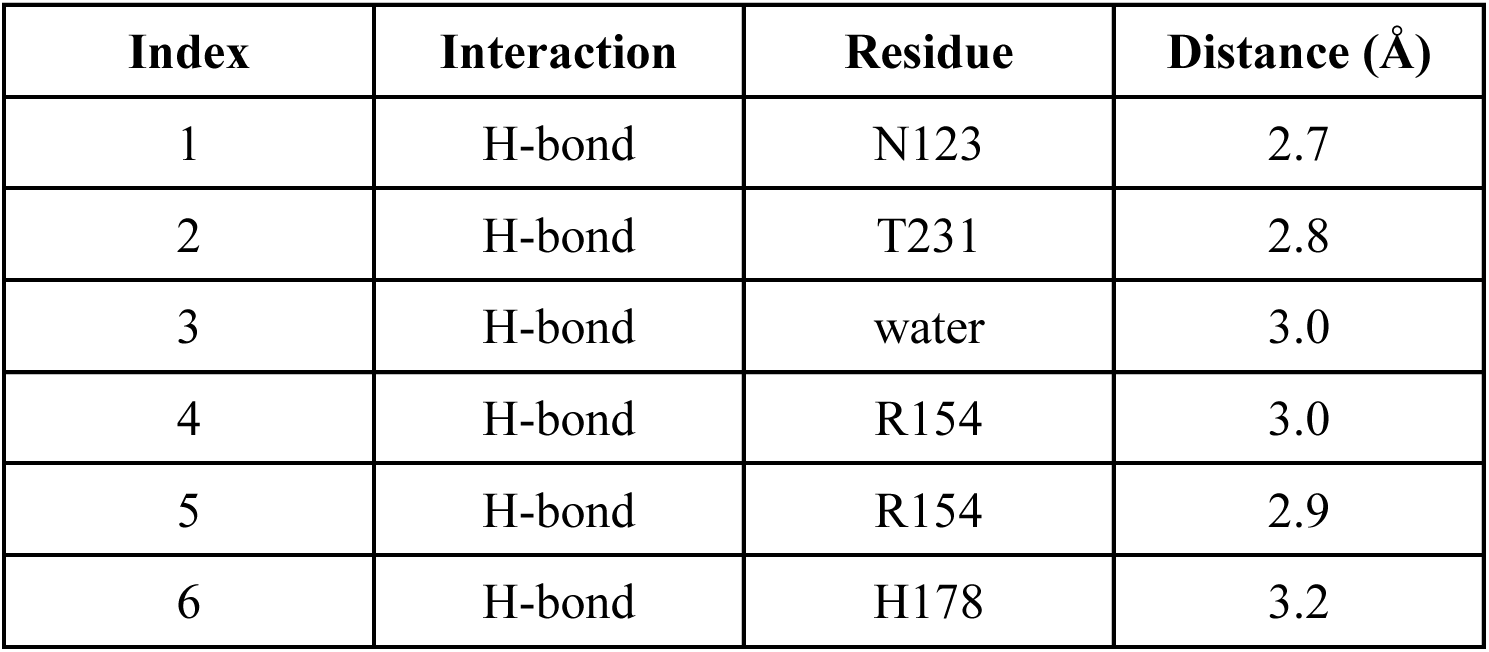
Interactions between LDH (chain A) and OXA.

| Index | Interaction | Residue | Distance (Å) |
| --- | --- | --- | --- |
| 1 | H-bond | N123 | 2.7 |
| 2 | H-bond | T231 | 2.8 |
| 3 | H-bond | water | 3.0 |
| 4 | H-bond | R154 | 3.0 |
| 5 | H-bond | R154 | 2.9 |
| 6 | H-bond | H178 | 3.2 |

**Table S4:** Interactions between LDH and FBP.

| Index | Interaction | Residue | Chain | Distance (Å) |
| --- | --- | --- | --- | --- |
| 1 | H-bond | water | - | 2.8 |
| 2 | H-bond | R156 | C | 3.1 x 2<br>3.2 |
| 3 | H-bond | H171 | C | 3.2 x 2<br>3.3<br>3.4 |
| 4 | H-bond | N169 | C | 2.5 |
| 5 | H-bond | Q168 | C | 2.6 |
| 6 | H-bond | H171 | A | 3.2 x 2<br>3.5 x 2 |
| 7 | H-bond | R156 | A | 3.2 x 2<br>3.4 |
| 8 | H-bond | N169 | A | 2.6 |

**Figure S1:**
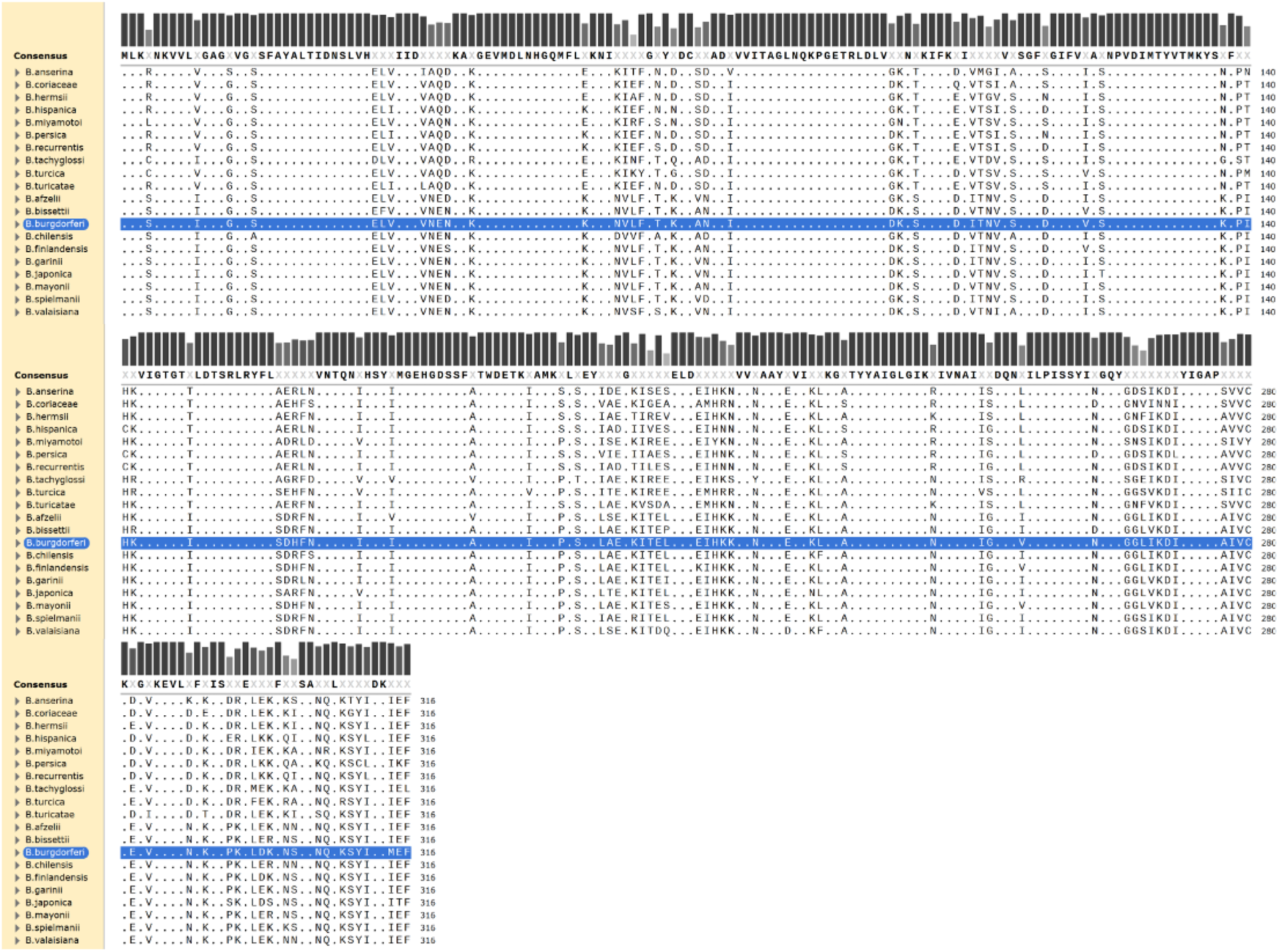
Sequence alignment of LDH across *Borrelia* species. LDH protein sequences for the *Borreliales* order (n = 20) were retrieved from AnnoTree (*1*), MSA generated via ClustalOmega as part of the MPI bioinformatics toolkit (*2, 3*) and visualized with SnapGene Viewer (SnapGene software (www.snapgene.com)). Sequence conservation is shown as gray bars, and a consensus sequence generated with a > 95% conserved threshold. Amino acids that are identical to the consensus sequence are indicated with a “**.”** and those that differ are denoted with their single-letter code. BbLDH sequence is highlighted for visualization purposes.

**Figure S2.**
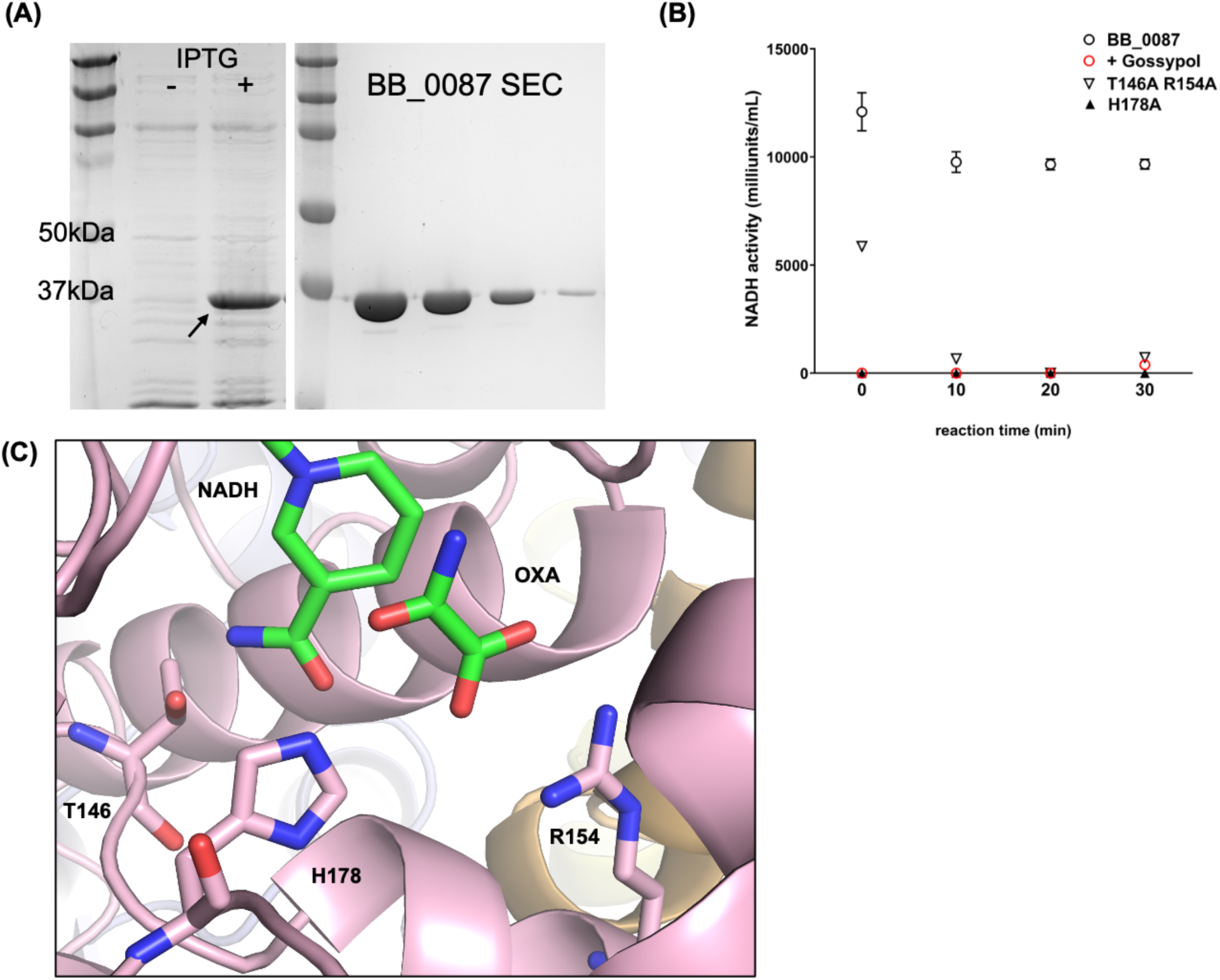
Expression and Purification of recombinant BB_0087. **(A)** Full length *bb_0087* sequence was cloned into a modified pET100/D-TOPO expression vector containing 10 x His and 1 x FLAG tag at the N-terminal and transformed into *E. coli* Rosetta-gami (DE3) expression cell line for induction. Upon the addition of 1 mM IPTG, induction of BB_0087 was observed around 37 kDa. Arrow indicates the induced protein. Purification of BB_0087 was performed using HiTrap His column followed by size exclusion chromatography column (SEC) to obtain homogenous protein. **(B)** BB_0087 has LDH activity. The LDH activity of purified BB_0087 was assessed using a Sigma Aldrich Lactate Dehydrogenase Assay Kit. NADH activity was detected using 50 ng of purified recombinant protein which was completely abolished upon treatment with 100 µM of gossypol. Mutation of conserved key residues (T146A R154A and H178A) found in LDH enzymes abolished the NADH activity of BB_0087. **(C)** Structure and location of conserved residues Thr-146, Arg-154 and His-178 in the NADH/OXA binding site of BbLDH. Figure was prepared using chain A, however, the location of all residues were invariant in all four chains of the BbLDH tetramer.

**Figure S3.**
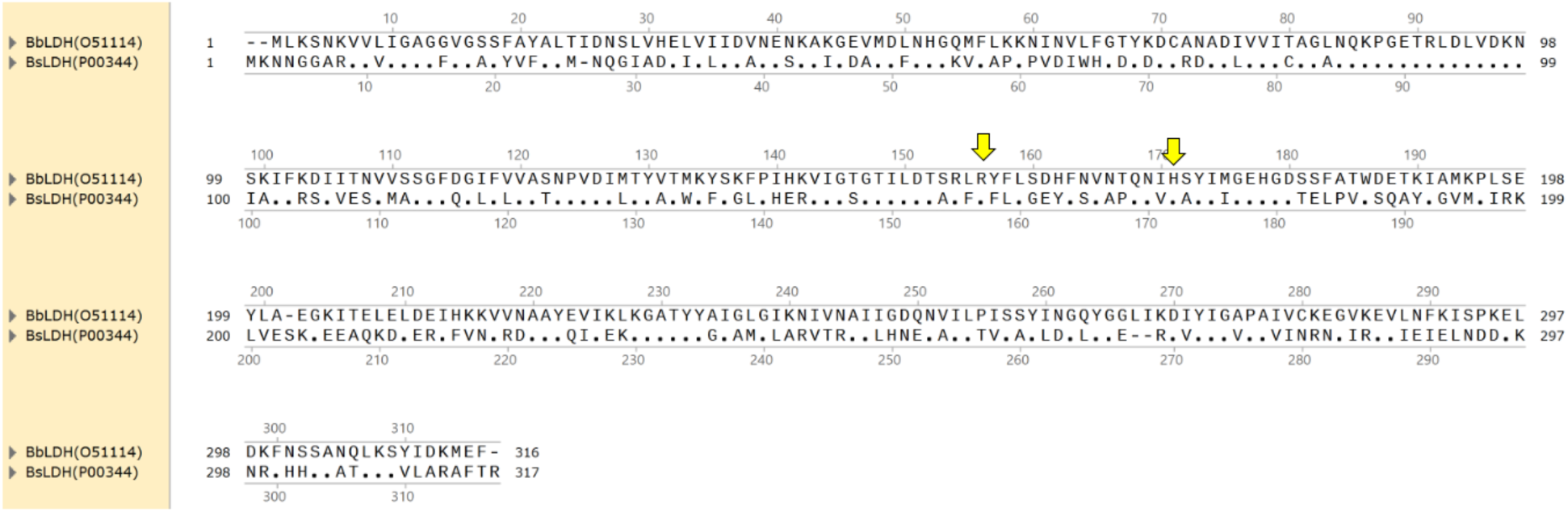
Sequence alignment between BbLDH (Uniprot ID: 051114) and BsLDH (Uniprot ID: P00344). Identical residues are indicated by a “.” and different residues are denoted with their one-letter residue code (for Bs). Conserved FBP binding residues are marked with a yellow arrow.

**Figure S4.**
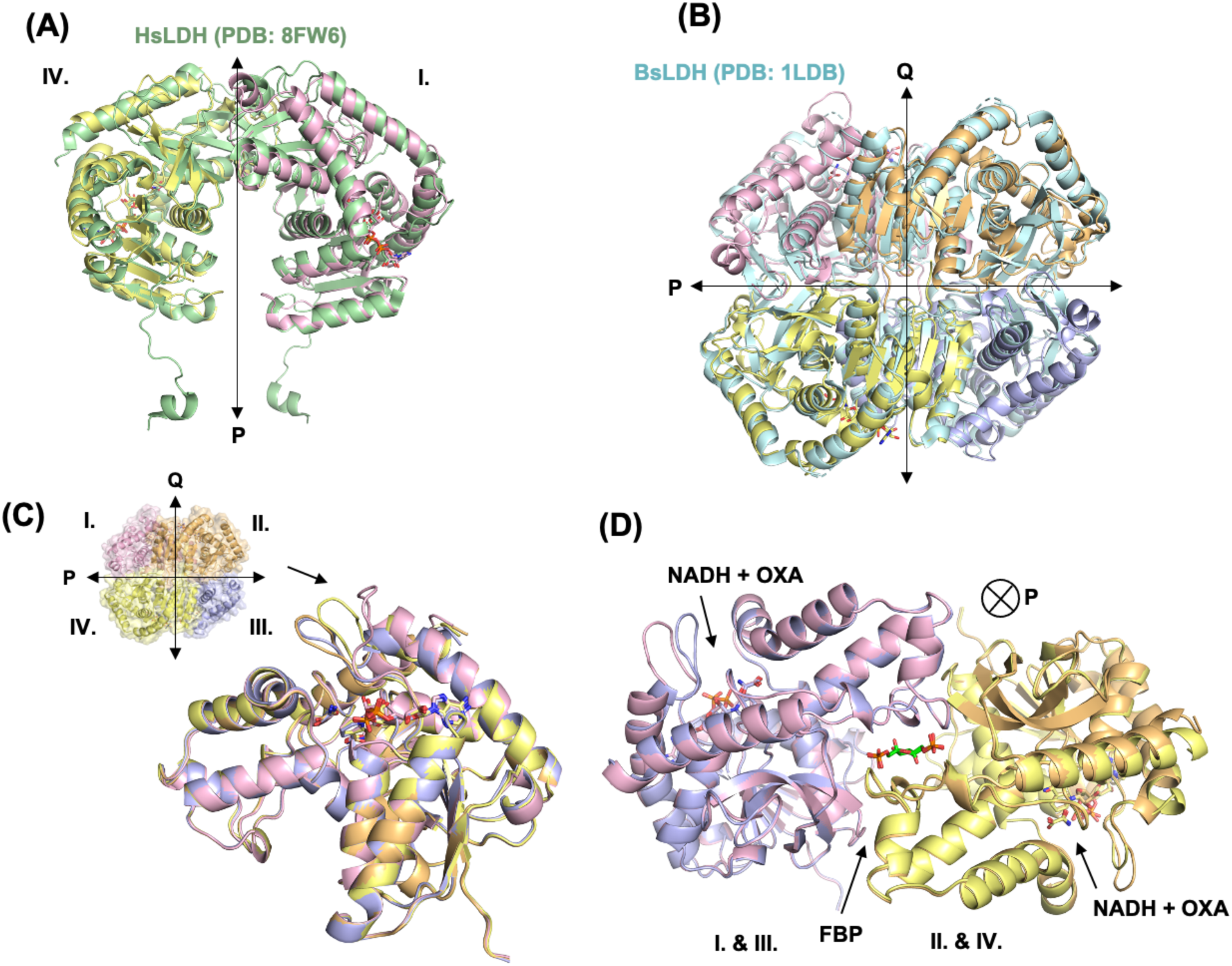
Structural comparison of BbLDH. **(A)** Superimposition of BbLDH dimer with HsLDH dimer (PDB 8FW6, RMSD = 1.394). **(B)** Superimposition of BbLDH and BsLDH tetramers (PDB 1LDB, RMSD = 17.270). **(C)** Structural comparison of the NADH and oxamate binding sites in each BbLDH monomer within the tetramer (RMSD range: 0.15-0.2). The major structural difference between all monomers is the relative position of the loop (Q85-R91) adjacent to the NADH/OXA binding site (black arrow). **(D)** Structural comparison between the two FBP binding sites per BbLDH tetramer (1 FBP per dimer).

**Figure S5.**
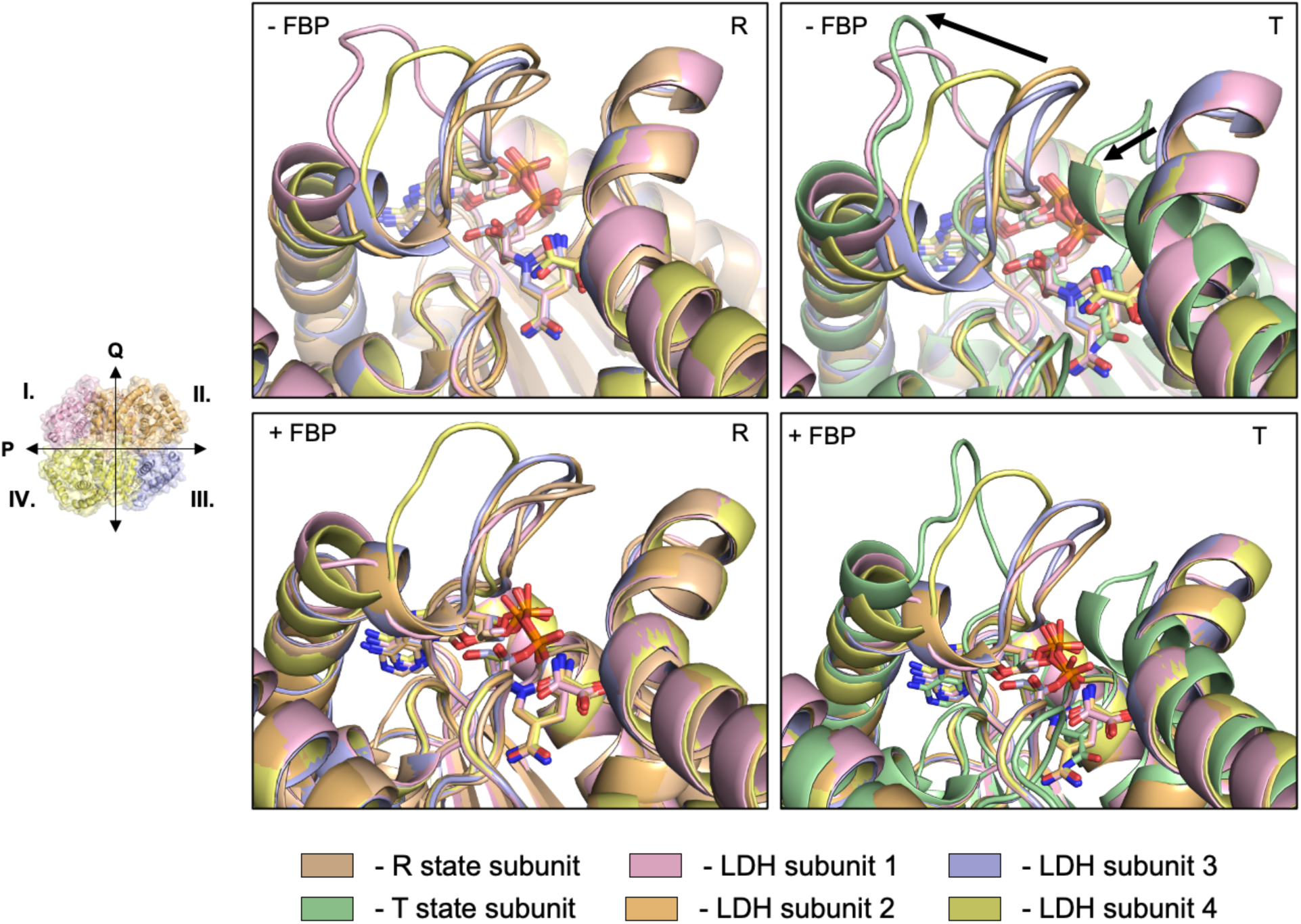
Structural comparison of BbLDH subunits to known R- and T-state conformations. Structural superimposition of LDH subunits from the BbLDH tetramer without (top) and with (bottom) FBP. In the left two panels *Bifidobacterium longum subsp. longum* (Bl, PDB: 1LTH) LDH subunit in the R-state (tan) is superimposed with the four BbLDH subunits, whereas in the right two panels the *B. longum* T-state subunit (green) is superimposed with the BbLDH subunits. Helix and loop motions that distinguish the T and the R states are shown with arrows on the top right.

**Figure S6.**
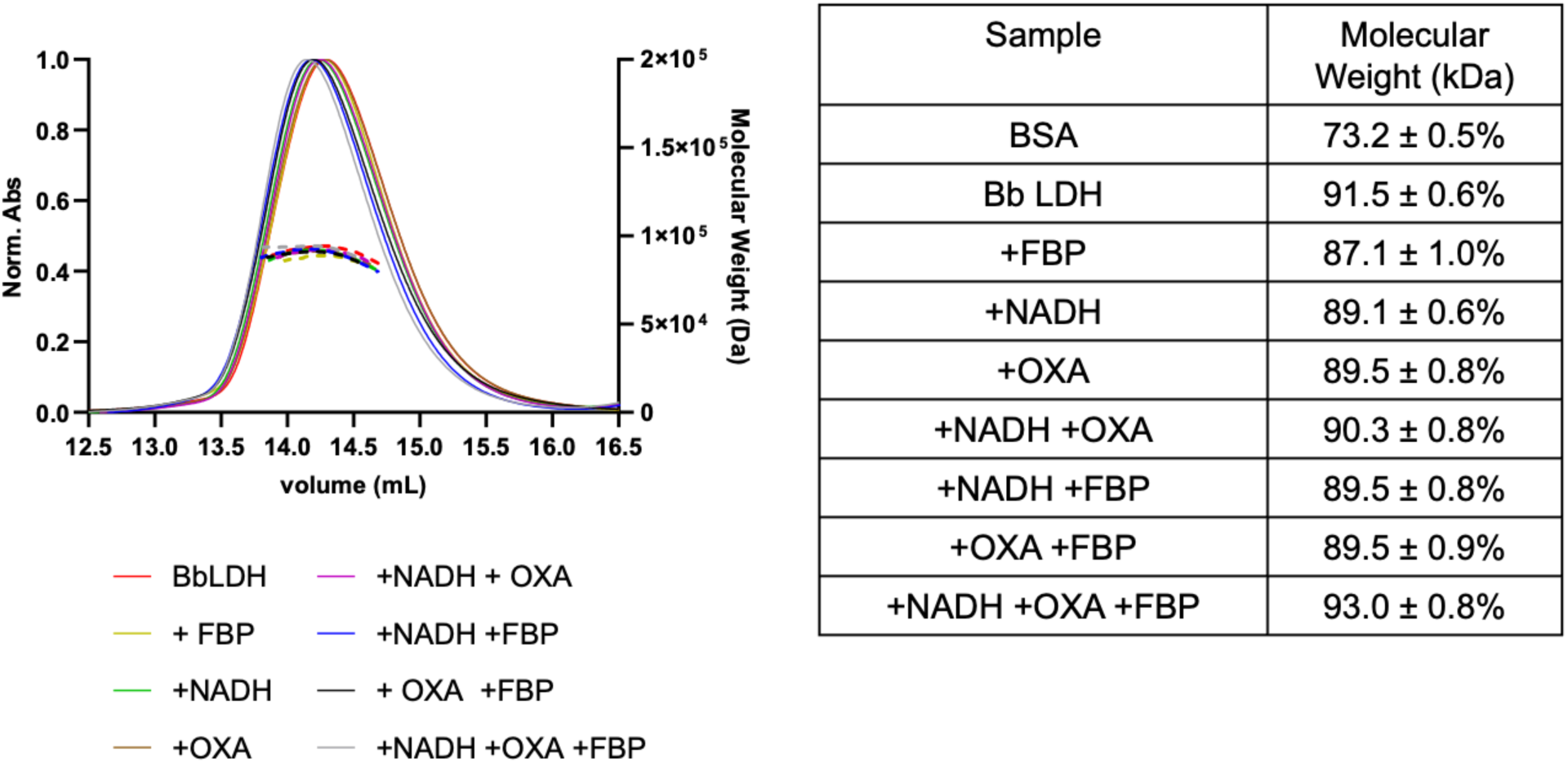
SEC-MALS of BbLDH with and without NADH, oxamate, and FBP. (left) SEC-MALS trace of all protein samples analyzed and (right) table of the molecular weights of each sample.

**Figure S7.**
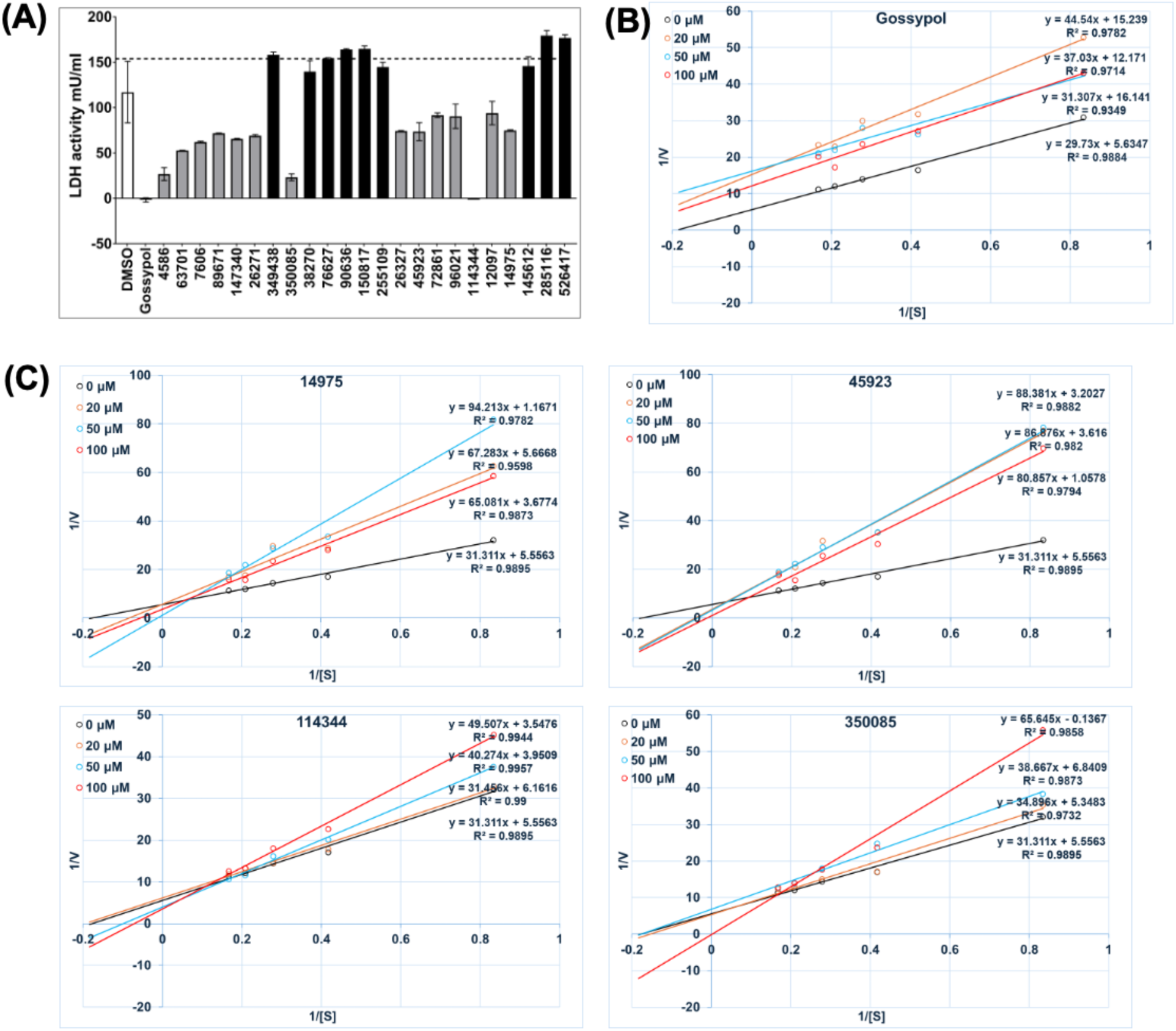
**(A)** An example of one set of screening results. DMSO was included as a vehicle control and gossypol as a positive control. Three hits (75% reduction in LDH activity as a cutoff) were identified: 350085 (270.28 Da), 45923 (216.18 Da), and 114344 (276.28 Da). Lineweaver–Burk plots of **(B)** gossypol and **(C)** 14975, 45923, 114344, and 350085 tested in this study.

**Figure S8.**
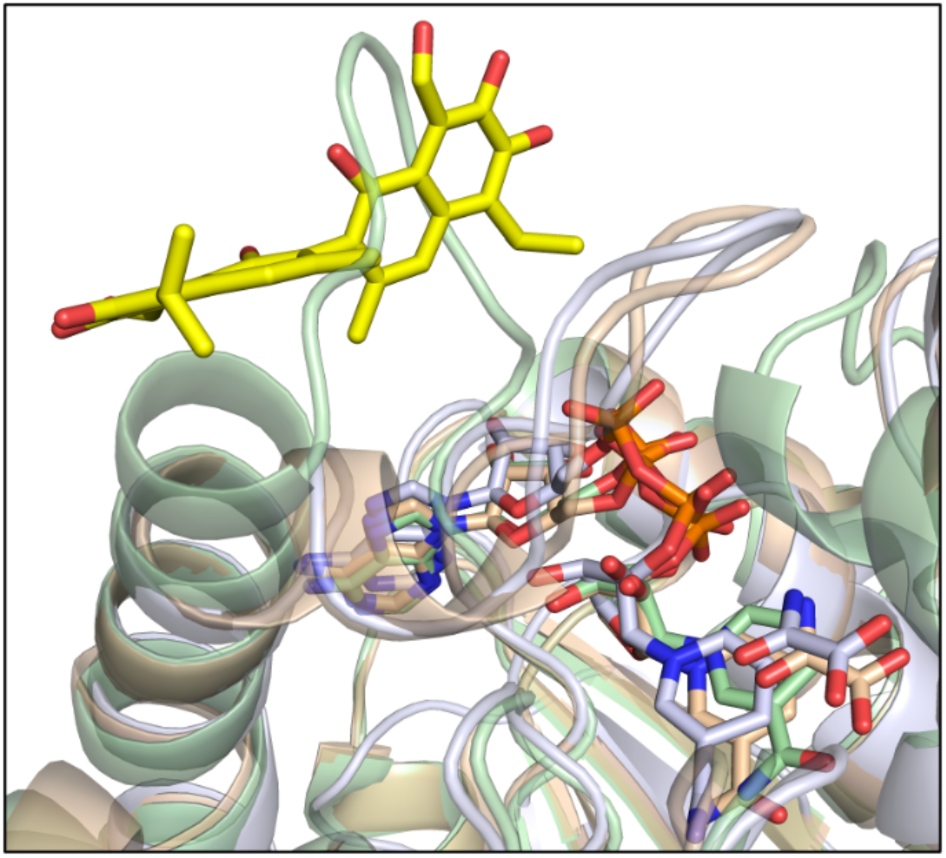
Gossypol docking pose with LDH model (light blue) with R- (tan) and T-state (green) BlLDH monomers (PDB: 1LTH).

**Figure S9.**
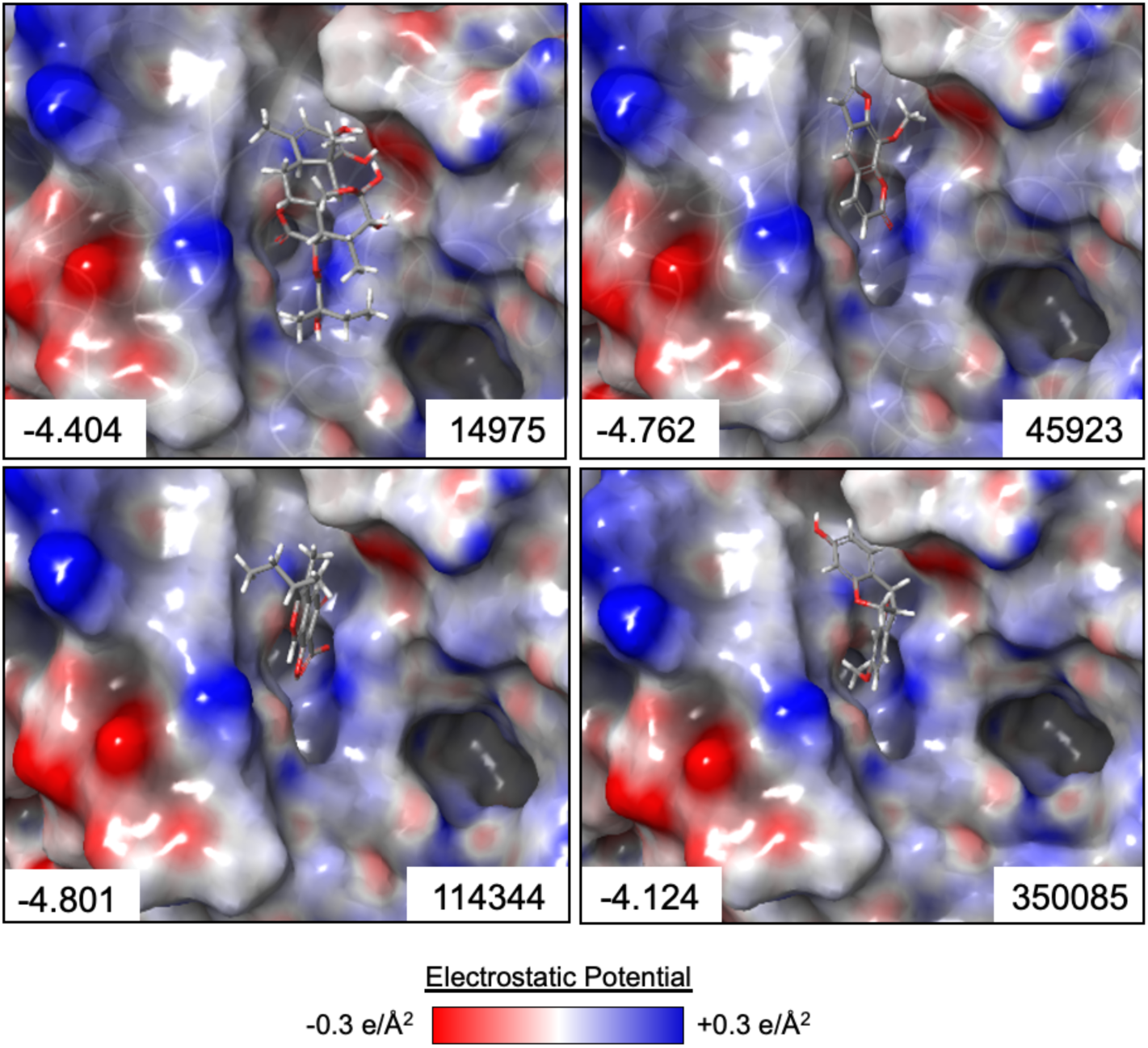
Top docking poses of 14975, 45923, 114344 and 350085. Top docking pose of each inhibitor to the apo BbLDH model. For each pose, the molecular surface of BbLDH is colored according to is electrostatic potential and the docking score denoted in the bottom right corner. (B)-(E) Ligand interaction diagram showing the hydrogen bonding interactions between each inhibitor and BbLDH.

**Figure S10.**
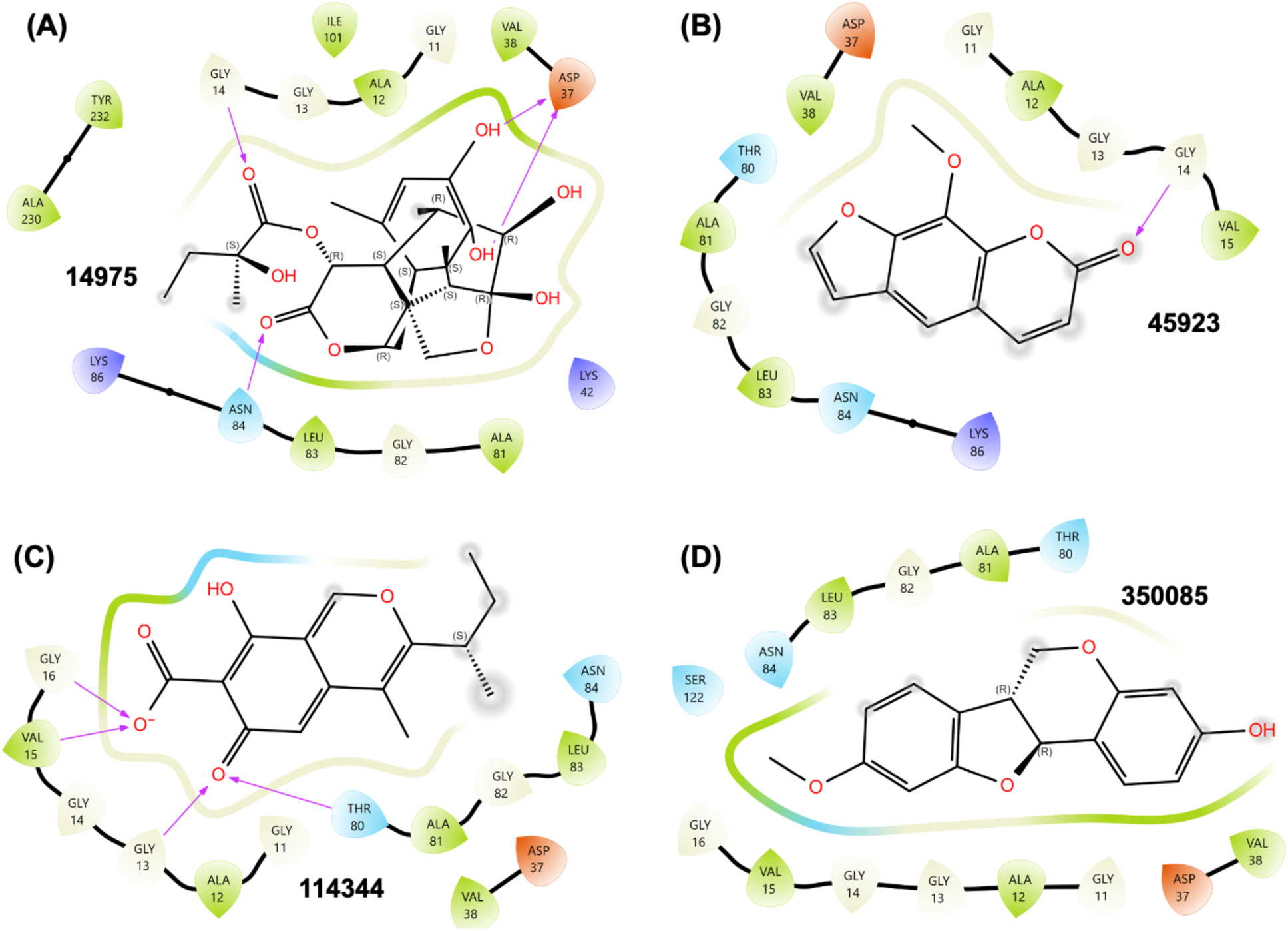
Interactions predicted by docking analysis. (A)-(D) Ligand interaction diagram showing the hydrogen bonding interactions between each inhibitor and BbLDH.

